# *Rickettsia parkeri* Hijacks Tick Hemocytes to Manipulate Cellular and Humoral Transcriptional Responses

**DOI:** 10.1101/2022.11.09.515877

**Authors:** Abdulsalam Adegoke, Jose M.C. Ribeiro, Sidney Brown, Ryan C. Smith, Shahid Karim

## Abstract

Blood-feeding arthropods rely on robust cellular and humoral immunity to control pathogen invasion and replication. Tick hemocytes produce factors that can facilitate or suppress microbial infection and pathogenesis. Despite the importance of hemocytes in regulating microbial infection, understanding of their basic biology and molecular mechanisms remains limited. Here we combined histomorphology and functional analysis to identify five distinct phagocytic and non-phagocytic hemocyte populations circulating within the Gulf Coast tick *Amblyomma maculatum*. Depletion of phagocytic hemocytes using clodronate liposomes revealed their function in eliminating bacterial infection. We provide the first direct evidence that an intracellular tick-borne pathogen, *Rickettsia parkeri*, infects phagocytic hemocytes in *Am. maculatum* to modify tick cellular immune responses. A massive RNA-seq dataset generated from hemocytes isolated from uninfected and *R. parkeri*-infected partially blood-fed ticks generated ∼40,000 differentially regulated transcripts, >11,000 of which were immune genes. Silencing two differentially regulated phagocytic immune marker genes (*nimrod B2* and *eater*) significantly reduced hemocyte phagocytosis. Together, these findings represent a significant step forward in understanding how hemocytes regulate microbial homeostasis and vector competence.

## 1. Introduction

Ticks are major vectors for bacterial, viral, and protozoan pathogens of human and veterinary importance. The Gulf Coast tick *Amblyomma* (*Am*.) *maculatum* is a competent vector of the spotted fever group *Rickettsia parkeri*, an obligate intracellular bacterium that causes an eschar-like lesions at the site of tick attachment (1, 2). Vector competence is influenced by the ability of a pathogen to establish infection, replicate, and disseminate across vector tissues, processes counteracted by the arthropod’s innate immune system. Similar to in vertebrates, the tick immune system has both cellular and humoral arms. The cellular arm is represented by hemocytes, which are professional immune cells equivalent to vertebrate leukocytes. Following microbial infection, tick hemocytes execute cell-mediated responses including phagocytosis, encapsulation, and nodulation, clearing microbes from the system (3–6). The humoral arm of the tick immune system produces soluble effector molecules that activate the complement pathway, prophenoloxidase pathway, and melanization cascade and produce reactive oxygen and nitrogen species (7–11). Hemocytes also secrete effectors that eventually activate humoral responses (12, 13).

Tick hemocytes have historically been classified into prohemocytes, plasmatocytes, granulocytes, spherulocytes, and oenocytoids (14) based on their quantity, size, shape, nuclear-cytoplasmic ratio, and presence of inclusion bodies. Recent ultrastructural studies have narrowed the classification to prohemocytes, granulocytes I, and granulocytes II (15-16), and functional characterization of tick hemocytes established the existence of core vertebrate hemocyte functions. Both hard and soft tick hemocytes use phagocytosis as the primary defense mechanism (3, 17–19), especially in the granulocyte and plasmatocyte subsets (8, 17, 20-21). Hemocytes can phagocytose tick-transmitted pathogens, such as hemocyte engulfment of *Borrelia* spirochetes, in a process described as coiling phagocytosis (22). *Ixodes scapularis* hemocytes were also reported to be infected with *Anaplasma phagocytophilum*, a requirement for subsequent salivary gland infection (23). The release of effector molecules complements hemocyte-mediated responses as part of the humoral defense response and include several pathogen recognition molecules such as lectins, antimicrobial peptides (AMPs), and thioester-containing proteins (24–26). However, the molecular mechanisms underlying tick hemocyte-pathogen interactions remain largely uncharacterized.

Here we used a conservative immunofluorescence and morphological approach to classify hemocyte subpopulations in *Am. maculatum*. Through *in vivo* phagocytosis of fluorescent beads, we functionally differentiate phagocytic from non-phagocytic hemocytes. For the first time, we demonstrate depletion of phagocytic hemocyte subsets using clodronate liposomes (CLD) in a tick species, thereby defining a role for phagocytic hemocytes in innate immune responses against bacterial challenge. Next-generation transcriptome analysis of uninfected and *R. parkeri-*infected hemolymph samples reveals molecular changes associated with pathogen recognition, immune pathway activation, and hemocyte production, amongst others. Using RNA interference, we characterize two previously unreported molecular markers of hemocyte phagocytosis in ticks. We also show that *R. parkeri* infects phagocytic hemocytes, thereby possibly playing a vital role in the systemic dissemination of *R. parkeri* to other tissues.

## 2. Materials and Methods

### 2.1 Ethics statement

All animal experiments were performed in accordance with the Guide for the Care and Use of Laboratory Animals of the National Institutes of Health, USA. Protocols for tick blood-feeding were approved by the University of Southern Mississippi’s Institutional Animal Care and Use Committee (USMIACUC protocols #15101501.3 and 17101206.2).

### 2.2 Tick maintenance and rearing

*Am. maculatum* ticks were maintained at the University of Southern Mississippi following previously established protocols for hard ticks (27). Established laboratory colonies of *Rickettsia parkeri-*infected ticks were generated from questing ticks. Unfed adult ticks were collected using the drag-cloth method during the summer months of 2019 from Mississippi Sandhill Crane, National Wildlife Refuge, Gautier, Mississippi (https://www.fws.gov/refuge/mississippi_sandhill_crane/). *R. parkeri-*free *Am. maculatum* ticks were obtained from the tick rearing facility at Texas A&M (TAMU, College Station, TX, USA) tick-rearing facility. These ticks were fed on cattle, which clears *R. parkeri* from tick tissues across feeding stages.

### 2.3 Hemolymph perfusion and hemocyte quantification

Before hemolymph collection, unfed or partially fed ticks were cleaned in a 10% sodium hypochlorite solution for 5 minutes, followed by a 10-minute wash in 70% ethanol and cleaning using double distilled water. Hemolymph was collected from ticks as previously described (28) using a freshly prepared modified citrate-EDTA anticoagulant buffer (vol/vol 60% Schneider’s *Drosophila* medium and 70% 5 mM EDTA in 1X PBS) (29) and kept on ice until needed. A modified perfusion method was used to collect hemolymph from unfed ticks. Briefly, ticks were placed with their ventral side facing downwards on double-sided tape mounted on a petri dish, which was then placed on ice for approximately 20 minutes to stimulate hemolymph flow within the tick. 0.5 µL of anticoagulant solution was injected into ticks between the basis capituli and scutum. Injected ticks were kept recovering at room temperature for 20 minutes before hemolymph perfusion. 1-2 injections were made between the ridges of the festoon using a 33G removable needle (Hamilton Company, Franklin, MA, USA). Immediately, 5 µL of anticoagulant solution was injected from the basis capituli and the exiting hemolymph was collected in a 1.5 mL microcentrifuge tube. Perfused hemolymph was centrifuged at 500 rpm for 3 minutes at 4°C to pellet the hemocytes, and the supernatant was removed. Hemocyte pellets were resuspended in a fresh anticoagulant buffer and kept on ice until needed to allow for the collection of hemolymph from unfed ticks. Total hemocytes were quantified using the trypan blue exclusion method (Invitrogen, Thermo Fisher Scientific, Waltham, MA, USA). Briefly, perfused hemolymph was mixed with 0.4% trypan blue, and 10 μL of the mixture was pipetted into a cell counting chamber. Hemocytes were quantified using a Countess automated cell counter (Invitrogen, Thermo Fisher Scientific Waltham, MA, USA). The number of hemocytes was estimated by placing perfused hemolymph onto the grooves of an improved hemocytometer chamber (Bright-Line, Hausser, Scientific Horsham, PA, USA), and hemocytes were differentiated morphologically as described previously (30,31).

### 2.4 Chemical depletion of phagocytic hemocytes

Phagocytic cells were depleted as previously described with slight modifications (32-33). To deplete circulating phagocytic hemocytes from the tick’s hemolymph, clodronate liposomes (CLD) or control liposomes (LP) (Standard Macrophage Depletion Kit, Encapsula Nano Sciences LLC, Brentwood, TN, USA) were injected into the hemolymph using a 2.5 μL 600 series Hamilton microliter syringe connected to a 33G removable needle (Hamilton Company). The optimal concentration of CLD and LP necessary for depletion with no adverse effect on survival was initially determined by injecting CLD and LP (stock, 1:2, 1:5, and PBS) in 1X PBS into groups of 15 ticks. These ticks were then monitored for survival over eight days beginning 24 h post-injection. Subsequent depletion experiments were performed using 1:5 dilution of CLD and LP in 1X PBS.

### 2.5 Bacterial challenge and survival analysis

*Escherichia coli* (*E. coli*) strain DH5-alpha and *Staphylococcus aureus* (*S. aureus*) strain RN4220 were grown overnight at 37□ in LB and TSA media, respectively. Overnight cultures were carefully removed, centrifuged, and adjusted to a concentration of OD_600_ = 0.5 for *E. coli* and OD_600_ = 0.1 for *S. aureus* in 1X PBS. *R. parkeri* were cultured as previously described (34). Frozen stocks of cell-free *R. parkeri* were revived by infecting Vero cells. After the cells were revived and replicated, the concentration of *R. parkeri* cells was determined using the plaque assay. Cells were isolated from Vero cells by lysis using sonication (BioRuptor™ Pico, Denville, NJ, USA) for 5 min in cycles of 30 s on and 30 s off at 4□. After sonication, the suspension was centrifuged at 1000 x g for 5 min at 4□ to pellet cell debris, and the supernatant was passed through a 0.22 µM syringe filter (Fisher Scientific, Grand Island, NY, USA). Cell-free *R. parkeri* was stored in SPG medium on ice until ready for use. Ticks were challenged with 0.2 µL bacteria 24 h post-CLD or LP injection. LPS and heat-killed *S. aureus* were used as positive controls for bacterial injection. The injection of sterile 1X PBS was used as a negative and injection control. Ticks were maintained and constantly monitored every 24 h for signs of mortality.

### 2.6 In vivo phagocytosis assay

The phagocytic function of tick hemocytes was assessed by injecting fluorescent-conjugated carboxylated beads as previously reported (33, 35) with slight modifications to adapt for use in ticks. Briefly, ticks were injected with 2% (vol/vol) 0.2 µL of yellow-green Carboxylated-Modified Microspheres (Thermo Fisher Scientific, Waltham, MA, USA) diluted in anticoagulant buffer and allowed to recover in incubators for 4 h at 22°C and 95% relative humidity (RH). Subsequently, hemolymph was perfused using an anticoagulant solution (vol/vol 60% Schneider’s *Drosophila* medium and 70% 5 mM EDTA in 1X PBS). Perfused hemolymph was allowed to adhere on a glass microscope slide for 1 h at room temperature. Hemocytes were fixed in 4% paraformaldehyde (PFA) in 1X PBS for an additional hour. Fixed hemocytes were washed with 1X PBS. Hemocytes were incubated with 20 μM Hoechst 33342 (Thermo Fisher Scientific) diluted in 1X PBS for 1 h at RT, after which slides were washed three times in 1X PBS and allowed to dry. Slides were mounted on a coverslip in 10 μL Fluoromount-G mounting medium (SouthernBiotech, Birmingham, AL, USA).

### 2.7 In-vivo EdU incorporation assay

We estimated hemocyte differentiation by visualizing and quantifying the synthesis of new DNA *in vivo* based on 5-ethynyl-2′-deoxyuridine (EdU) incorporation and subsequent detection using the Click-iT EdU Alexa Fluor 647 kit (Invitrogen, Grand Island, NY, USA) as previously described (36,37), with the only exception that 0.5 µL of 20 mmol l−1 EdU in anticoagulant buffer was injected into ticks. Ticks were allowed to recover in incubators for 4 h at 22°C and 95% RH. Following recovery, hemolymph was perfused from ticks and allowed to attach on a microscope glass slide for 1 h at 4°C. Hemocytes were subsequently fixed with 4% paraformaldehyde diluted in 1X PBS for 30 min at RT, washed three times with 3% bovine serum albumin (BSA) in 1X PBS, permeabilized for 30 min with 0.5% Triton-X in PBS at RT, followed by another wash step with 3% BSA in 1X PBS. Hemocyte slides were subsequently incubated in the dark with the Click-iT reaction cocktail for 30 min at RT according to the manufacturer’s instructions, followed by a wash step with 3% BSA in 1X PBS. Hemocyte slides were subsequently incubated with 20 μM Hoechst 33342 (Thermo Fisher Scientific) diluted in 1X PBS for 1 h at RT, after which slides were washed three times in 1X PBS and allowed to dry before mounting on a microscope glass slide by adding 10 μL Fluoromount-G mounting medium (SouthernBiotech).

### 2.8 Hemocyte staining

Tick hemolymph was perfused into an anticoagulant buffer and allowed to adhere to a glass coverslip in a humid chamber at 4°C for 1 h. Without washing, hemocytes were fixed by adding 4% PFA solution in 1X PBS for an additional 1 h at RT. After fixation, cells were washed three times with 1X PBS and permeabilized with 0.1% Triton X-100 for 1 h at RT. Without washing, hemocytes were blocked in 1% BSA solution in 0.1% Triton X-100 for an additional 1 h at RT. Excess blocking solution was washed with 1X PBS. Hemocytes were incubated with 1U phalloidin (Alexa Fluor™ 488 Phalloidin, Molecular Probes, Thermo Fisher Scientific) and 20 μM Hoechst 33342 (Molecular Probes, Thermo Fisher Scientific) diluted in 1X PBS for 1 h at RT, after which slides were washed in 1X PBS and allowed to dry before mounting on a microscope glass slide by adding 10 μL Fluoromount-G mounting medium (SouthernBiotech).

### 2.9 RNA extraction, cDNA synthesis, and qRT-PCR

Hemolymph was collected from 10 individual ticks as described above and pooled, and an anticoagulant buffer was added to a total volume of 250 μL. RNA was extracted using the Trizol-chloroform separation and isopropanol precipitation method with slight modifications. Following the initial chloroform separation of RNA into the aqueous phase, a second separation was performed by adding a 1:1 volume of chloroform to the aqueous phase, centrifuging at maximum speed, and the transparent upper phase was used to proceed with isopropanol precipitation. Second, an ethanol wash of the RNA pellet was carried out twice to help completely remove the isopropanol carryover. The RNA pellet was air dried and resuspended in 30 μL of nuclease-free water, concentration and quality checked, and stored at −80°C until use. Complementary DNA synthesis and qRT-PCR were conducted as previously described (38). Sequences of gene-specific primers designed to amplify cDNA fragments are listed in **Supplementary Table S1**. Transcriptional gene expression was normalized against the *Am. maculatum* β*-actin* gene. The synthesized cDNA was used to measure mRNA levels by qRT-PCR using the CFX96 PCR Detection System (Bio-Rad Inc., Hercules, CA, USA) described previously (38–40).

### 2.10 Double-stranded RNA (dsRNA) synthesis and delivery

Double-stranded RNA from *nimrod B2* and *eater* transcripts was synthesized for gene silencing and microinjected into unfed adult female *Am. maculatum* ticks as previously described (38–42). Before injection, dsRNA targeting each gene was diluted to a working concentration of 1 μg/μL in nuclease-free water. Double-stranded DNA from the green fluorescent protein (*Gfp*) gene was synthesized and injected as an irrelevant control.

### 2.11 Illumina sequencing

RNA samples from uninfected and *R. parkeri-*infected *Am. maculatum* hemolymph were sent for sequencing by Novogene (China). Briefly, partially blood-fed (∼50 mg, slow blood feeding phase and ∼200 mg, start of fast feeding phase) ticks were selected for RNA sequencing. Hemolymph was collected from 120 partially blood-fed ticks during the slow-feeding (∼50 mg) and fast-feeding phases (∼200 mg). Three biological replicates of hemolymph from the slow-feeding and fast-feeding phases were included in each sample of the *R. parkeri-*infected or uninfected group, i.e., a total of 12 samples. Hemolymph from ten partially-fed ticks was combined for each biological replicate. Hemolymph RNA was extracted as described above. RNA libraries were constructed from hemolymph RNA from six uninfected and six *R. parkeri-* infected ticks using the NEBNext Ultra™ RNA library Prep Kit (New England Biolabs, Ipswich, MA, USA). RNA library preparation and sequencing were conducted by Novogene Co., Ltd. (Beijing, China).

### 2.12 Bioinformatics analysis

Raw reads were stripped of contaminating primers, and bases with qual values <20 were trimmed. Clean reads were assembled using the Abyss (43) and Trinity (44) assemblers. Resulting contigs were re-assembled with a blastn and cap3 assembler (45) pipeline as described previously (46). Coding sequences were extracted based on blastx results derived from several database matches, including a subset of the non-redundant NCBI protein database containing tick and other invertebrate sequences, as well as the Swiss-Prot and Gene Ontology (GO) databases. All open reading frames larger than 200 nucleotides were extracted, and those matching known proteins or with a signal peptide were retained. The resulting peptide and coding sequences were mapped to a hyperlinked spreadsheet including blastp and rpsblast matches to several databases and an indication of the signal peptide (47), transmembrane domains (48), and O-galactosylation sites (49). edgeR was used in ancova mode to detect statistically significant differentially-expressed genes according to feeding or infection status (50). edgeR inputted the read matrix for genes with at least one library expressing an FPKM (fragments per thousand nucleotides per million reads) equal to or larger than 10. For heat map visualization of CDS temporal expression, Z scores of the FPKM values were used. All deduced coding sequences and their reads are available for browsing with hyperlinks to several databases (**Supplementary Table S2**).

### 2.13 RNA-seq and differential gene expression analysis

As previously described, differentially expressed genes from edgeR analysis were analyzed using the iDEP (integrated Differential Expression and Pathway analysis) online tools (51). The expression matrix representing read counts of differentially-expressed genes and the gene IDs were uploaded to the iDEP user interface and used for data exploration.

### 2.14 Immunofluorescence of R. parkeri

Hemolymph from unfed or partially fed ticks was perfused onto a microscope coverslip, and hemocytes were allowed to adhere for 1 h at RT. Hemocytes were fixed in 4% PFA (4% in PBS; J19943-K2, Thermo Fisher Scientific) for 30 min at RT. Coverslips containing hemocytes were permeabilized with 0.1% Triton X-100 for 30 min at RT. For non-permeabilized coverslips, 1X PBS was added to the hemocytes for 30 min at RT. This step was followed by three times washing with 1X PBS. Non-specific proteins were blocked with 1% BSA solution in PBS for 1 h, followed by primary incubation with mouse anti-*Rickettsia* M14-13 (generously provided by T. Hackstadt, NIH/NIAID Rocky Mountain Laboratories (52,53)), and rabbit anti-*Sca-2* (generous gift from Matthew D. Welch, UC Berkeley). A no primary antibody control sample (negative control) was prepared in parallel. *R. parkeri* was detected using goat anti-rabbit Alexa Fluor 568 and goat anti-mouse Alexa Fluor 568 (1:500 in 1% BSA; Invitrogen, Thermo Fisher Scientific). Samples were washed three times in 1X PBS to remove free antibodies. Hemocytes were incubated with 20 μM Hoechst 33342 (Molecular Probes, Thermo Fisher Scientific) diluted in 1X PBS for 1 h at RT, after which slides were washed three times in 1X PBS and allowed to dry before mounting on a microscope glass slide by adding 10 μL Fluoromount-G mounting medium (SouthernBiotech).

### 2.15 Imaging acquisition

Confocal images were acquired with a Leica STELLARIS STED (Leica Microsystems, Wetzlar, Germany) confocal microscope using either a 40X, 63X, or 100X objective (zoom factor 3-5; numerical aperture of 1). Images were obtained using both sequential acquisition and variable z-stacks. The 405 UV laser was used to acquire the DAPI channel, while the tunable white light laser (WLL) was used to capture the Alexa-Fluor channel. A z-stack of the images consisting of 150-250 slices was compiled for all images captured, and the proprietary Leica built-in post-processing plugin was used for deconvolution and to carry out lightning processing. All images were exported as acquired and compiled in PowerPoint software.

### 2.16 Data availability

Raw fastq files were deposited in the Sequence Read Archives (SRA) of the National Center Biotechnology Information (NCBI) under bioproject PRJNA878782 and biosamples SAMN30755417 and SAMN30755418. Deduced coding sequences and their translations were deposited in the Transcriptome Shotgun Assembly database DDBJ/EMBL/GenBank under the accessions GKCB01000001-GKCB01011171.

## 3. Results

### 3.1 Discrimination of hemocyte types

Light microscopic examination of direct hemolymph smears revealed a heterogeneous hemocyte population. There were two distinct small and large cell populations, the latter comprising cells with varying cytoplasmic contents, nuclear shape, and cytoplasm size (**Supplementary Fig. S1A and B**). The position of the nucleus was variable: certain hemocytes possessed large, centrally placed nuclei occupying most of the cytoplasmic space, while in some cells the nucleus was peripheral and binucleated. Variable granulation was also observed in the cytoplasm of some hemocyte types.

However, the resolution of light microscopy limited our discrimination of hemocyte subsets. We therefore assessed whether we could further classify hemocytes using commonly used fluorescent markers. Perfused hemolymph was stained with wheat germ agglutinin (WGA), Vybrant CM-Dil (a lipophilic cell membrane stain), and Hoechst 33342 (a nuclear stain). WGA discriminated hemocyte populations with varying degrees of binding intensity, indicating a potential difference in hemocyte function based on their lectin binding activity. By contrast, all hemocytes were positive for CM-Dil (**Supplementary Fig. S1C**). To further differentiate between hemocyte subtypes, we co-stained with phalloidin (an actin stain) and DAPI (a nuclear stain). Five distinct hemocyte types were identified based on shape, actin projections, and nuclear-cytoplasmic ratio: (i) granulocytes were relatively large and had multiple actin projections; (ii) plasmatocytes were pyriform with a centrally placed nucleus; (iii) spherulocytes possessed a peripherally placed nucleus; (iv) prohemocytes were characterized by a relatively high nuclear to cytoplasmic ratio; and (v) oenocytoids had a smaller nuclear to cytoplasmic ratio (**Fig. 1A**).

The total hemocyte population differed significantly between partially fed and unfed female ticks, with hemocyte numbers increasing after the blood meal (**Supplementary Fig. S1D**). Feeding also affected the hemocyte subtype distribution in male and female ticks: feeding significantly increased granulocyte numbers in females (**Supplementary Fig. S1E**) and significantly decreased the spherulocyte population in both male and female ticks (**Supplementary Fig. S1F**). Plasmatocyte numbers were lower in blood-fed females than unfed females but increased in males on feeding (**Supplementary Fig. S1G**). There was a trend to oenocytoids numbers increasing following feeding in both males and females (**Supplementary Fig. S1G**), but prohemocytes were absent in both male and female ticks following feeding (**Supplementary Fig. S1I**).

Hemocyte populations were histomorphologically similar in male and female ticks, so we next examined functional differences between male and female hemocytes by assessing their phagocytic abilities. Phagocytic hemocytes were more common in female ticks than in male ticks in an *in vivo* phagocytosis assay of yellow-green FluoSpheres (**Fig. 1B and C**). Taken together, these data demonstrate that *Am. maculatum* hemocyte heterogeneity might influence hemocyte function.

**Figure 1.**
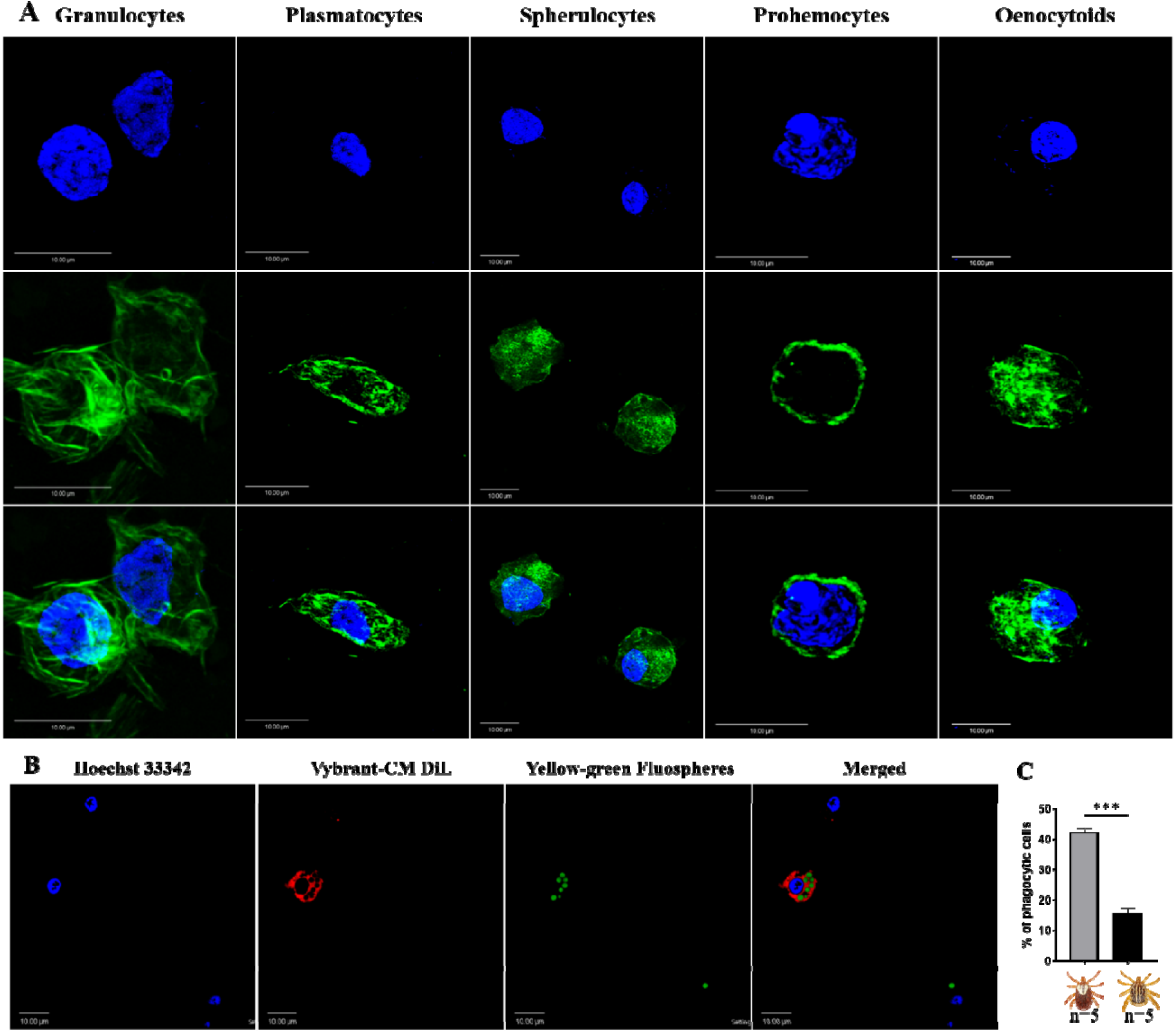
Confocal microscopy images of *Am. maculatum* hemocytes stained with phalloidin (green) and Hoechst 33342 (blue). Hemocytes were subtyped based on nuclear size and location and cytoplasmic projections **(A)**. Granulocytes are relatively large and have multiple actin projections. Plasmatocytes are pyriform and have a centrally placed nucleus. Spherulocyte possess a peripherally placed nucleus. Prohemocytes are characterized by a relatively large nuclear to cytoplasmic ratio and oenocytoids by a smaller nuclear to cytoplasmic ratio. Functional assessment of hemocyte phagocytosis of green FluoSpheres **(B)** and the proportion of phagocytic hemocytes in male and female ticks **(C)**. Quantitative data were analyzed using unpaired *t-*tests in GraphPad Prism v8.4.1. \**P* < 0.05, \*\**P* < 0.01, \*\*\**P* < 0.001, \*\*\*\**P* < 0.0001. Scale bar = 10 μm.

### 3.2 Clodronate liposomes deplete and impair phagocytic hemocyte functions

In the absence of definitive molecular markers of hemocyte subtypes in ticks, there is a need for alternative tools to study the role of hemocytes in cellular immunity and vector competence. To this end, clodronate liposomes (CLD), a pharmacological agent that specifically targets and depletes professional phagocytes via apoptosis (32-33, 54-55), was used to deplete phagocytic hemocytes in tick hemolymph. CLD exclusively targets cells with phagocytic abilities (56). Upon phagocytosis, the liposome is degraded by the lysosome, which releases toxic clodronate to promote apoptosis (**Fig. 2A**).

**Figure 2.**
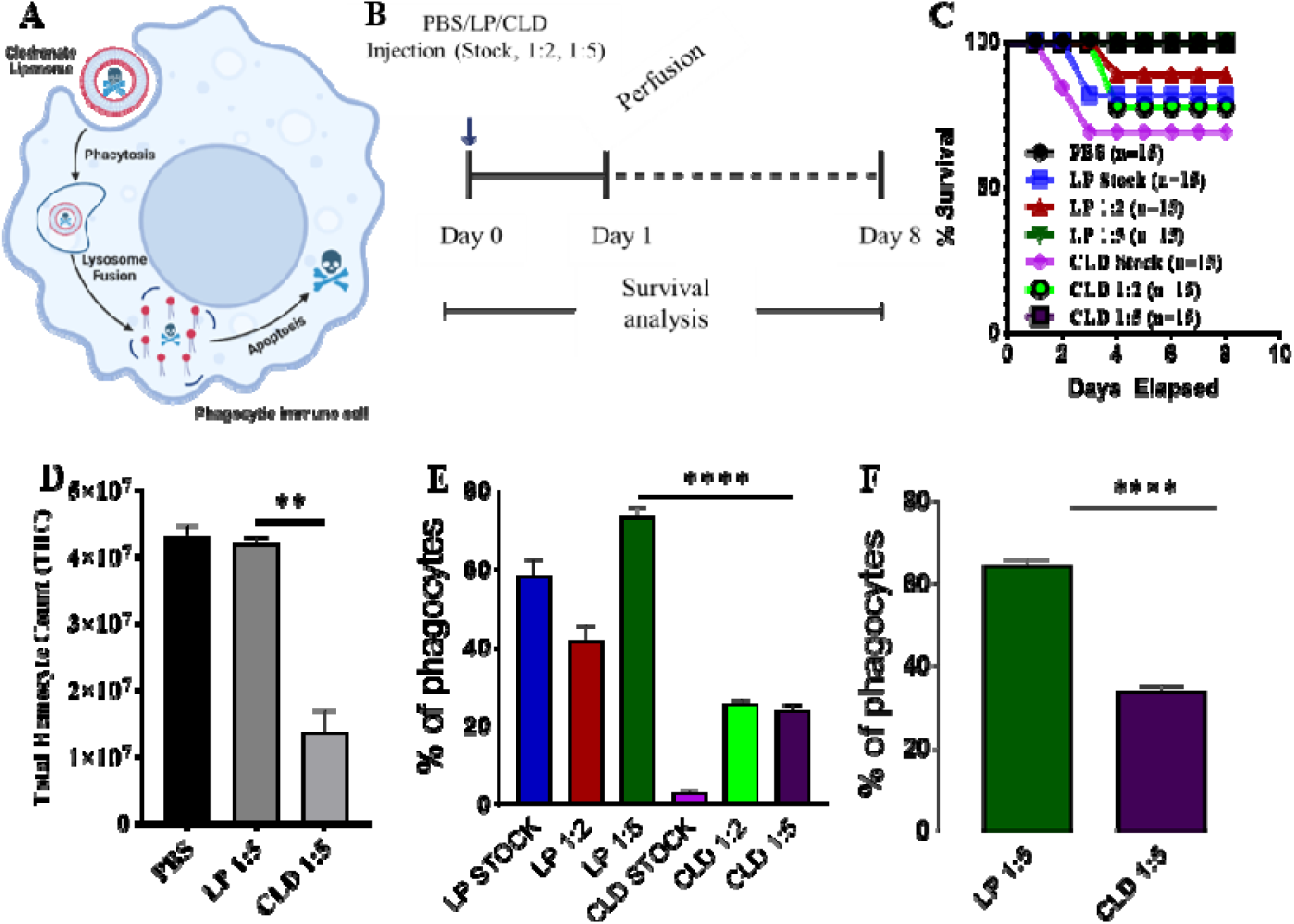
Clodronate depletion of phagocytic tick hemocytes and validation of phagocyte depletion. Mechanism of clodronate liposome-induced depletion of professional phagocytes **(A)**. Schematic showing the optimization of clodronate and liposome concentrations to deplete phagocytic hemocytes **(B)**. Tick survival was evaluated following injection of clodronate (CLD) and control liposomes (LP) at different concentrations (stock, 1:2, 1:5 in 1X PBS), with 1X PBS used as control **(C)**. Hemolymph was perfused 24 h post-CLD or LP injection (unfed status) to assess the effect of depletion on total hemocyte count **(D)** and proportion of phagocytic hemocytes **(E)**. The proportion of phagocytic hemocytes was also assessed in CLD- or LP-injected ticks 5-days post feeding **(F)**. Survival was checked each day for 8 days; 15 ticks were assigned to each treatment group. Significance was determined with the log-rank (Mantel-Cox) test using GraphPad Prism v8.4.1. Error bars represent ± SEM of five ticks. Quantitative data were analyzed using unpaired *t-*tests in GraphPad Prism v8.4.1. \**P* < 0.05, \*\**P* < 0.01, \*\*\**P* < 0.001, \*\*\*\**P* < 0.0001.

We first tested different clodronate and control liposome (LP) concentrations to determine their impact on tick survival (**Fig. 2B**). Injection of CLD or LP at a 1:5 dilution had no adverse impact on tick survival (**Fig. 2C**) but significantly reduced total hemocyte populations (**Fig. 2D**) due to reduced numbers of phagocytic hemocytes (**Fig. 2E**). To determine the effect of the blood meal on phagocytic hemocyte depletion, we injected unfed ticks with a 1:5 dilution of CLD or LP before a blood meal and quantified the phagocytic hemocyte population following partial feeding. The blood meal did not interfere with the ability of CLD to deplete phagocytic hemocytes (**Fig. 2F**).

To further assess the effect of CLD depletion on phagocyte function, ticks injected with CLD or LP were fed and *in vivo* hemocyte phagocytosis assayed with yellow-green FluoSpheres. Hemolymph of LP-injected ticks contained more hemocytes that engulfed one or two FluoSpheres than hemocytes from clodronate-depleted ticks (**Fig. 3A**). Hemocyte populations have previously been shown to increase following a blood meal due to cellular division (37), so we determined the impact of clodronate depletion on hemocyte DNA replication. Partially blood-fed CLD or LP-injected ticks were injected with EdU, and their hemocytes were assayed for EdU incorporation into hemocyte DNA. Significantly more EdU was incorporated into LP-injected tick hemocytes compared with CLD-injected ticks (**Fig. 3B**). Together, these data confirm effective chemical depletion of the phagocytic hemocyte population and show that CLD interferes with the abilities of hemocytes to undergo replication and differentiation.

**Figure 3.**
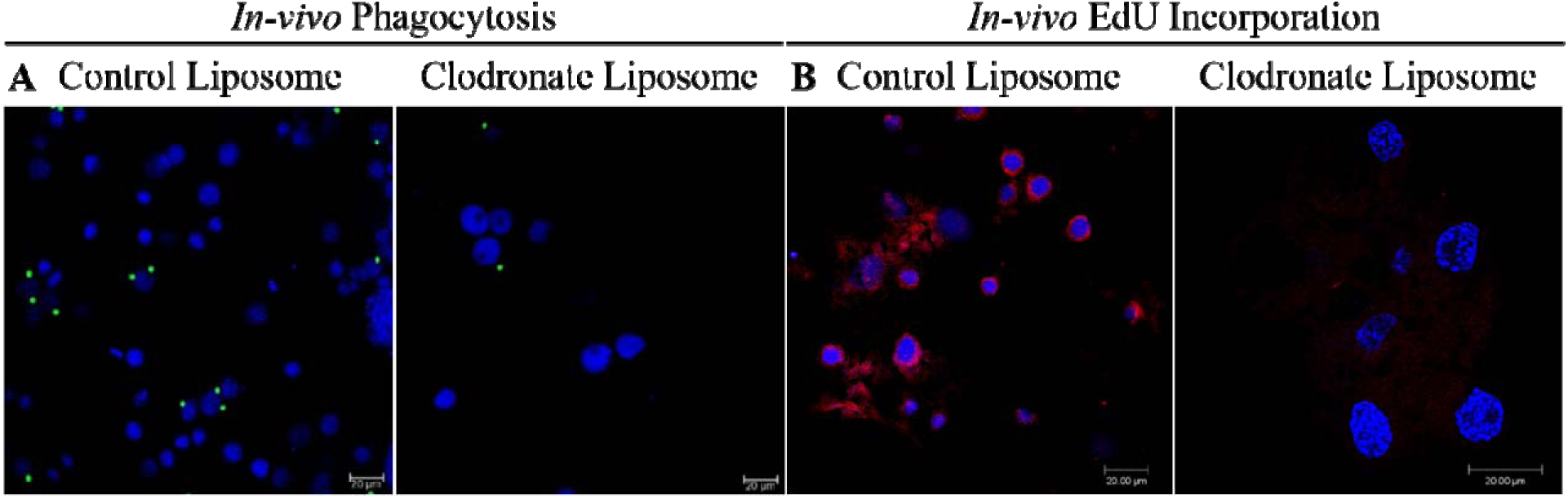
Clodronate liposomes deplete hemocyte functions. Clodronate liposomes impaired hemocyte phagocytosis **(A)** and interfered with EdU incorporation into hemocytes DNA **(B)**. Scale bar = 20 μm.

### 3.3 Depletion of phagocytic hemocytes impairs survival against bacterial challenge

Hemocytes are a vital defense mechanism against invading microbes in ticks, mosquitoes, and *Drosophila* (32, 33). We therefore attempted to determine how immunocompromised ticks would survive challenge with both Gram-positive and Gram-negative bacteria (**Fig. 4A**). Ticks were unaffected by injection with PBS (**Fig. 4B**). Phagocyte depletion impaired tick survival against Gram-negative *E. coli* (**Fig. 4C**), but Gram-positive *S. aureus* (live and heat-inactivated) significantly affected tick survival in both CLD and LP-treated groups (**Fig. 4D**). Similarly, phagocyte depletion significantly impaired survival against *Am. maculatum-*transmitted *R. parkeri* (**Fig. 4E**). Since phagocytic granulocytes act as scavengers of invading microbes (6,32,33), these data further support the hypothesis that phagocytic hemocytes are critical components of the immune response in ticks and maintain tick microbial homeostasis via cell-mediated immunity and interactions with pathogenic microbes (6, 57).

**Figure 4.**
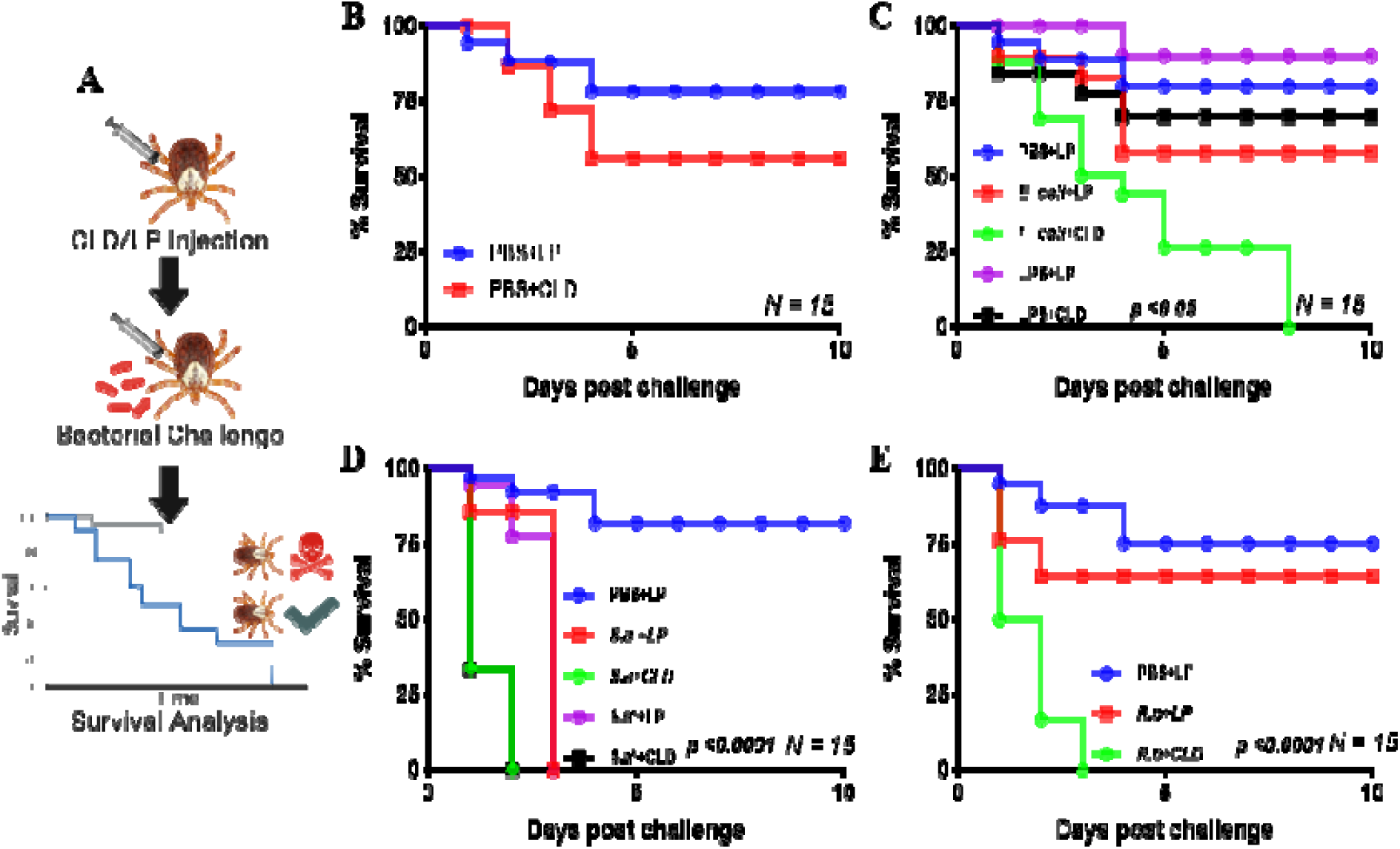
Depletion of phagocytic hemocytes impairs survival against bacterial challenges. Unfed female *Am. maculatum* were injected with either LP or CLD at 1:5 dilution and 24 h later challenged with bacteria or sterilely injured **(A)**. Tick survival was monitored every 24 h for 10 days to evaluate the effect of sterile injury **(B)**, *E. coli* **(C)**, *S. aureus* **(D)**, or *R. parkeri* **(E)** challenge. Data were analyzed with the log-rank (Mantel-Cox) in GraphPad Prism v8.4.1. *S*.*a: S. aureus, S*.*a*^*\**^: *heat-killed S. aureus, R*.*p: R. parkeri*.

### 3.4 R. parkeri can infect circulating hemocytes

The tick hemolymph contains a heterogeneous population of circulating hemocytes, as shown by ourselves (**Fig. 1A**) and others (4,7,10,16,31,58–60). *R. parkeri* acquired during a blood meal must circumvent both cellular and tissue barriers in the midgut to access hemolymph for systemic dissemination and subsequent transmission to a mammalian host. In the hemolymph, *R. parkeri* must either avoid, evade, or suppress hemocyte-mediated immune responses to successfully disseminate. *R. parkeri* is closely related to *Anaplasma* (*A*.) *phagocytophilum*, both existing as obligate intracellular pathogens and belonging to the same Rickettsiales order. Dissemination of *A. phagocytophilum* to the salivary gland in its tick vector is facilitated by direct hemocyte infection following midgut colonization by the bacteria (23). Since our understanding of how *R. parkeri* disseminates through the hemolymph from the midgut to other tissues is still limited, we asked whether *R. parkeri* can infect circulating hemocytes. Hemolymph from infected and unfed female *Am. maculatum* were incubated with primary antibodies targeting the outer membrane of *R. parkeri*, and *R. parkeri* actively entered the cytoplasm of hemocytes (**Fig. 5A**). In addition, intracytoplasmic infection was detected in the hemocytes of uninfected ticks previously injected or capillary-fed with GFP-expressing *R. parkeri* (**Fig. 5B and C**). While positive Sca-2 staining suggested the presence of *R. parkeri* in hemocytes, this could also have occurred through binding of *R. parkeri* to the surface of hemocytes arising from the immune response. However, detection of *R. parkeri* Sca-2 signal in both permeabilized and unpermeabilized hemocytes further confirmed active entry into hemocytes. These data argue that tick hemocytes - and potentially phagocytic hemocytes - are infected by *R. parkeri*, which might be important for its systemic dissemination. Similar findings had been reported with *A. phagocytophilum* and Zika virus infection of tick and mosquito hemocytes, respectively (61), further corroborating our observations.

**Figure 5.**
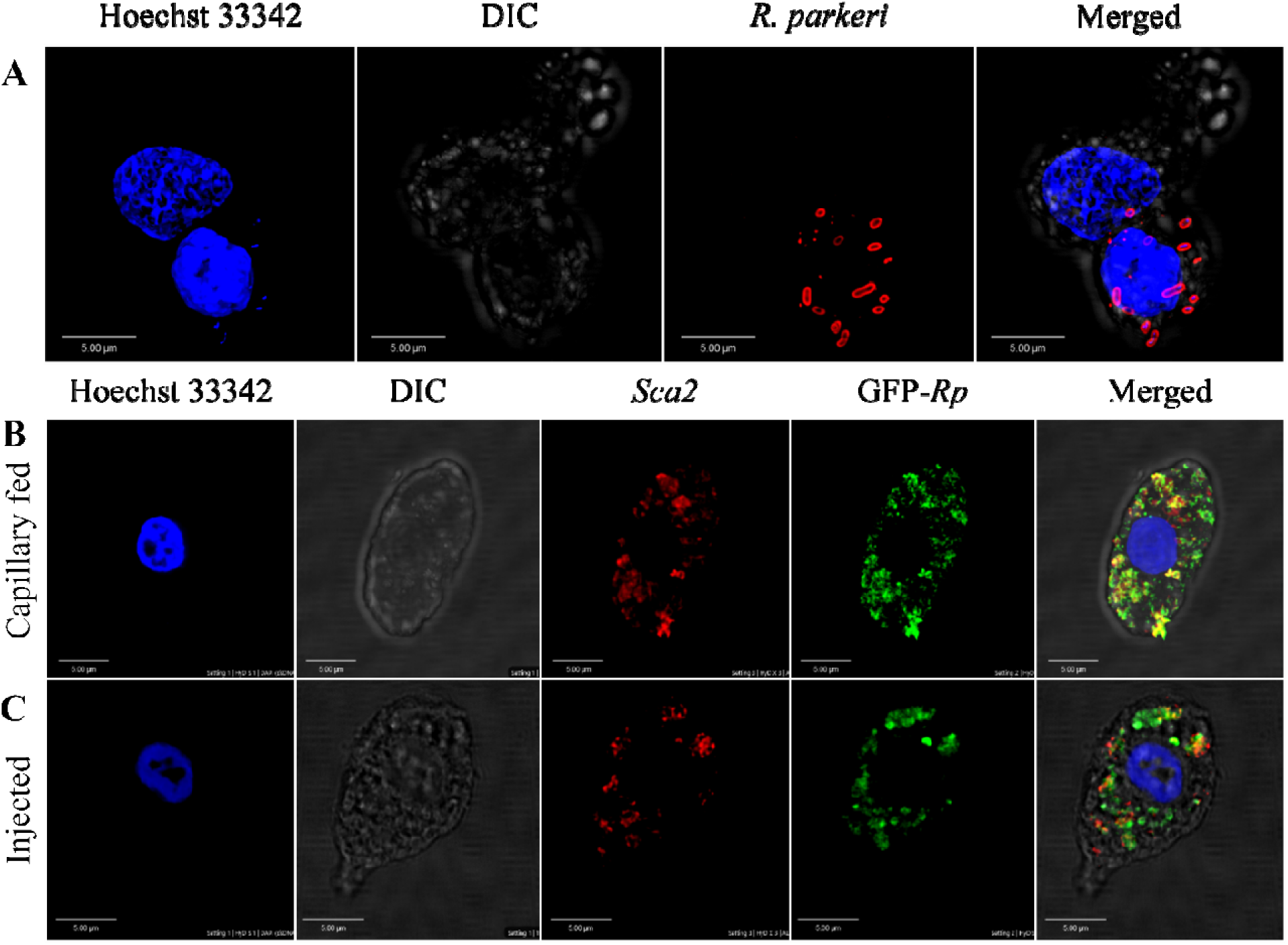
Confocal microscopy images of phagocytic hemocytes infected with *Rickettsia parkeri*. Representative confocal images of immunofluorescence staining for *R. parkeri* proteins showing hemocytes from natural and artificially infected ticks. **(A)** Immunolocalization of *R. parkeri* in hemocytes of naturally infected *Am. maculatum*. Hemocytes were incubated with primary antibodies targeting *R. parkeri* outer membrane protein (red) and Hoechst 33342 (blue). Infection of hemocytes with *R. parkeri* following **(B)** capillary feeding and **(C)** microinjection of GFP-expressing *R. parkeri* into uninfected *Am. maculatum*. Hemocytes were incubated with *R. parkeri* Sca-2 antibody (red) and Hoechst 33342 (blue). Scale bar = 5 μm.

### 3.5 RNA-seq reveals changes in hemocyte gene expression associated with R. parkeri infection

Identifying hemocyte-specific transcripts is crucial for the discovery of immune factors participating in hemocyte-mediated immune responses. While tick hemocyte transcripts have previously been characterized (62), the effect of pathogen infection on tick hemocyte gene expression is unknown. Here we generated and compared hemocyte transcripts from *R. parkeri*-infected (n = 6) and uninfected (n = 6) *Am. maculatum*. Assembly of the 305,276,990.3 reads from 12 libraries allowed us to identify 37,430 CDS, and database searching matched the CDS into 28 categories (**Table 1**). Secreted proteins accounted for 18.7% and 38.8% of the coding sequences and mapped reads, respectively, and included enzymes, protease inhibitors, lipocalins, and immune-related genes. Coding sequences classified with immune functions represented 0.86% and 2.45% of the total CDS and reads, respectively. More than 30% of coding sequences and 3.1% of total reads represented sequences of unknown function. Sorting of significantly expressed coding sequences with their respective reads identified 2,859 differentially expressed CDS between *R. parkeri-*infected and uninfected hemocytes (**Table 2**). Of these, 938 belonged to the secreted class, 46 to the immunity class, 568 were unknown, and 48 were associated with cytoskeletal functions. Forty bacterial-derived coding sequences were also significantly differentially expressed between *R. parkeri-*infected and uninfected hemocytes. In addition, we identified 39 coding sequences with functions as regulators of hematopoiesis, hemocyte differentiation, and immune functions. Of the 39, 14 (30.8%) were significantly differentially regulated between *R. parkeri-*infected and uninfected hemocytes (**Table 3**), and these genes include *transglutaminases, astakines, hemocytin, prokineticin, thymosin, eater, nimrod B2*, Runt-related transcription factor (*Runx*) and GATA-binding factor (*GATA*). We also identified coding sequences in the Toll and immune deficiency (IMD) immune pathways.

### 3.6 Regulators of hematopoiesis and hemocyte differentiation

#### 3.6.1 GATA factors and Runt domain-containing sequences

Seven genes regulating hemocyte production and differentiation were identified. Two transcription factors, *GATA* (AmHem-289270, AmHem-210536, AmHem-273849, AmHem-239125, AmHem-347209, AmHem-347211, AmHem-482569, AmHem-482570, and AmHem-470445) and *Runt* (AmHem-441656 and AmHem-441653) were differentially regulated between infected and uninfected hemocytes (**Supplementary Fig. S3A**). Seven of the nine GATA transcription factors and all the Runt domain-containing sequences were significantly upregulated in *R. parkeri-*infected hemocytes. GATA factors and Runt proteins have previously been shown to be critical in maintaining pluripotent hemocyte precursors in the hematopoietic organ (24,63).

#### 3.6.2 Astakines, β-thymosin, and transglutaminases

We identified one *astakine* (AmHem-345670) and three β*-thymosin* (AmHem-338118, AmHem-338117, and AmHem-338116) coding sequences differentially regulated upon *R. parkeri* infection (**Supplementary Fig. S3A**). *R. parkeri* led to a three-fold upregulation of the AmHem-345670 transcript (*astakine*) and downregulation of the three β*-thymosin* transcripts (AmHem-338118, 11-fold; AmHem-338117, 2-fold; and AmHem-338116, 2-fold). *Astakines* are ancient cytokines with conserved cysteine domains that share similar homology to vertebrate prokineticins (64,65). β*-thymosins* are small peptides involved in numerous cellular processes such as cellular migration, tissue repair and cell adhesion, proliferation, and differentiation in vertebrates. Their affinity for ATP-synthase is crucial to their function (66). Like β*-thymosins, transglutaminases* (TGases) are ubiquitously expressed and regulate many cellular processes such as cellular adhesion, cell migration, and the maintenance of the extracellular matrix. Six TGases (AmHem-380205, AmHem-552537, AmHem-482635, Amac-hemSigP-334835, AmHem-459192, and AmHem-174366) were significantly upregulated, except for AmHem-482635 and Amac-hemSigP-334835, which were downregulated on *R. parkeri* infection (**Supplementary Fig. S3A**).

#### 3.6.3 Laminin receptors and CLIP-domain serine proteases

Laminin receptors are a group of proteins with diverse biological functions, including cellular differentiation. They serve as binding partners with different homeostasis-associated proteins to maintain hemocyte homeostasis (67). The *CLIP*-domain serine protease (*CLIPsp*) is abundantly present in the hemolymph of insects and arthropods. Four transcripts with a Clip or disulfide knot domain (AmHem-164283, Amac-hemSigP-457530, AmHem-462558, and AmHem-462556) were differentially expressed in our dataset (**Supplementary Fig. S3A**): *R. parkeri* led to six-fold upregulation of AmHem-AmHem-164283, 14-fold upregulation of Amac-hemSigP-457539, and two-fold upregulation of AmHem-462558 and AmHem-462556. In invertebrates, these proteins play dual roles in innate immune responses and hematopoiesis, acting as binding partners of *toll*-like receptor *Spaetzle*, leading to downstream transcriptional activation of antimicrobial peptides. Similarly, they activate the prophenoloxidase (PPO) cascade necessary for melanization (68). A direct role has been described for *CLIPsp*-induced PPO maintenance of hematopoiesis (64).

### 3.7 Regulators of hemocyte-mediated cellular functions

Hemocyte-mediated cellular responses are an important component of the invertebrate innate immune system, and hemocyte functions, such as phagocytosis, are relatively conserved across invertebrate species. Several cell surface receptors are involved in the cellular immune response. In *Drosophila* and mosquitoes, the eater and nimrod transmembrane receptor families of proteins serve as phagocytosis receptors and scavenge bacteria for phagocytic killing. AmHem-270031 (*eater*) and AmHem-305744 (*nimrod B2*) were significantly downregulated (>9-fold) in *R. parkeri-*infected hemocytes (**Supplementary Fig. S3B**). Thioester-containing proteins (TEPs) are like the mammalian complement system and are involved in microbial opsonization prior to phagocytosis. We found three TEPs in our transcriptome data, with all transcripts (AmHem-349981, AmHem-459726, and AmHem-349977) containing an alpha-2-macroglobulin domain (**Supplementary Fig. S3B**). AmHem-349981 and AmHem-349977 were 5-fold upregulated, while AmHem-459726 was 3-fold downregulated in *R. parkeri-*infected hemocytes. Four alpha-2-macroglobulin (α2-macroglobulin) transcripts (AmHem-43749, AmHem-473966, AmHem-340857, and AmHem-241896), each consisting of the complement component region of the alpha-2-macroglobulin family, were also differentially expressed (**Supplementary Fig. S3B**). Three of the four α2-macroglobulin transcripts (AmHem-473966, AmHem-340857, and AmHem-241896) were >3-fold upregulated, while AmHem-43749 was 9-fold downregulated in *R. parkeri-*infected hemocytes. However, AmHem-43749 was only expressed in two of the six uninfected hemocyte groups.

AmHem-369012, AmHem-320301, AmHem-358999, AmHem-441843, AmHem-207626, and AmHem-289376 were transcripts containing secretory signal peptides with class F scavenger receptor domains, and all were significantly downregulated >6-fold in *R. parkeri-*infected hemocytes. Fourteen transcripts (Amac-hemSigP-444130, Amac-hemSigP-470263, AmHem-396331, AmHem-396326, AmHem-487866, AmHem-477539, AmHem-477540, AmHem-376633, AmHem-337488, AmHem-376632, Amac-hemSigP-396114, AmHem-199732, AmHem-295667, and Amac-hemSigP-336380) containing fibrinogen-related domains (FReDs) were significantly downregulated in *R. parkeri-*infected ticks (**Supplementary Fig. S3B**). FReD-containing proteins are involved in complement activation and phagocytosis in mammals (69), and several have been identified in invertebrates such as crabs (70), snails (71), mosquitoes (72,73), and ticks (62,74). Our data also showed significant downregulation (>10-fold) of two transcripts (AmHem-310057 and AmHem-310058) with a lectin C-type domain and mannose-binding activity. The binding activities of lectins make them suitable for pathogen recognition and are an important component of the immune response.

### 3.8 Regulators of the Toll pathway

The Toll pathway is highly conserved in both insects and other arthropod species. The peptidoglycan recognition receptor proteins (PGRPs) recognize lysine-type peptidoglycan on the cell wall of Gram-positive bacteria. In contrast, recognition of fungal β1-3-glucan occurs via the Gram-negative binding proteins (GNBPs) (75,76). This binding leads to translocation of nuclear factor kappa B (NF-κB) into the nucleus and subsequent upregulation of antimicrobial peptides. Eight PGRP transcripts (AmHem-345000, AmHem-459642, Amac-hemSigP-212567, Amac-hemSigP-370787, AmHem-140793, AmHem-429218, AmHem-310141, and AmHem-308618) were identified in our RNA-seq dataset. Seven of the PGRP transcripts were 3-10-fold downregulated following *R. parkeri* infection, while AmHem-429218 was 3-fold upregulated. AmHem-459642, Amac-hemSigP-370787, and AmHem-308618 are secreted, while AmHem-140793 is the only differentially expressed membrane-bound PGRP transcript in our dataset (**Supplementary Fig. S4**). Nine genes encoding Toll-related receptors have been reported in *Drosophila* (77), with some yet to be identified in the tick genome. All the components of the Toll pathway were detected and differentially regulated in our transcriptome data (**Supplementary Fig. S4A and B**) except for *GNBP, Tube*, and *Pelle*, the latter two gene products forming a heterodimer with MyD88 in *Drosophila* (77). Activation of *Spaetzle*, a ligand for the Toll receptor, is the rate-limiting step leading to activation of the Toll pathway. AmHem-476392 (6-fold upregulated) and AmHem-475327 (2-fold upregulated) were differentially expressed in *R. parkeri-*infected hemocytes (**Supplementary Fig. S4A**). Two Toll receptors with a leucine-rich repeat ribonuclease inhibitor domain, AmHem-352197 (4-fold upregulated) and AmHem-450801 (2-fold upregulated), were significantly expressed in our dataset (**Fig. 6A, Supplementary Fig. S4**). However, of the MyD88-Tube-Pelle heterotrimeric complex, only one *Myd88* transcript, AmHem-151635 (upregulated), was detected in our dataset. AmHem-417516 (4-fold upregulated), an ankyrin repeat and DHHC-type Zn-finger domain-containing protein encoding Cactus, a negative regulator of the Toll pathway that binds and prevents nuclear translocation of two Rel proteins, Dorsal and Diff (78), was significantly expressed in our data. We also identified a homolog of *Dorsal*, AmHem-31466 (2-fold upregulated), and *Diff*, Amac-hemSigP-143049 (7-fold downregulated), both containing a Rel homology domain (RHD) of RelA and RelB respectively (**Fig. 6A, Supplementary Fig. S4**). Nuclear translocation of Dorsal and Diff regulates AMP expression, especially the defensin and drosomycin family of AMPs. From our data, ten *Defensin* transcripts (Amac-hemSigP-382382, AmHem-345595, AmHem-286925, Amac-hemSigP-347294, Amac-hemSigP-433534, AmHem-296786, Amac-hemSigP-284010, AmHem-314023, AmHem-205330, and AmHem-482396) were significantly downregulated in *R. parkeri-*infected hemocytes (**Fig. 6A, Supplementary Fig. S4**).

**Figure 6.**
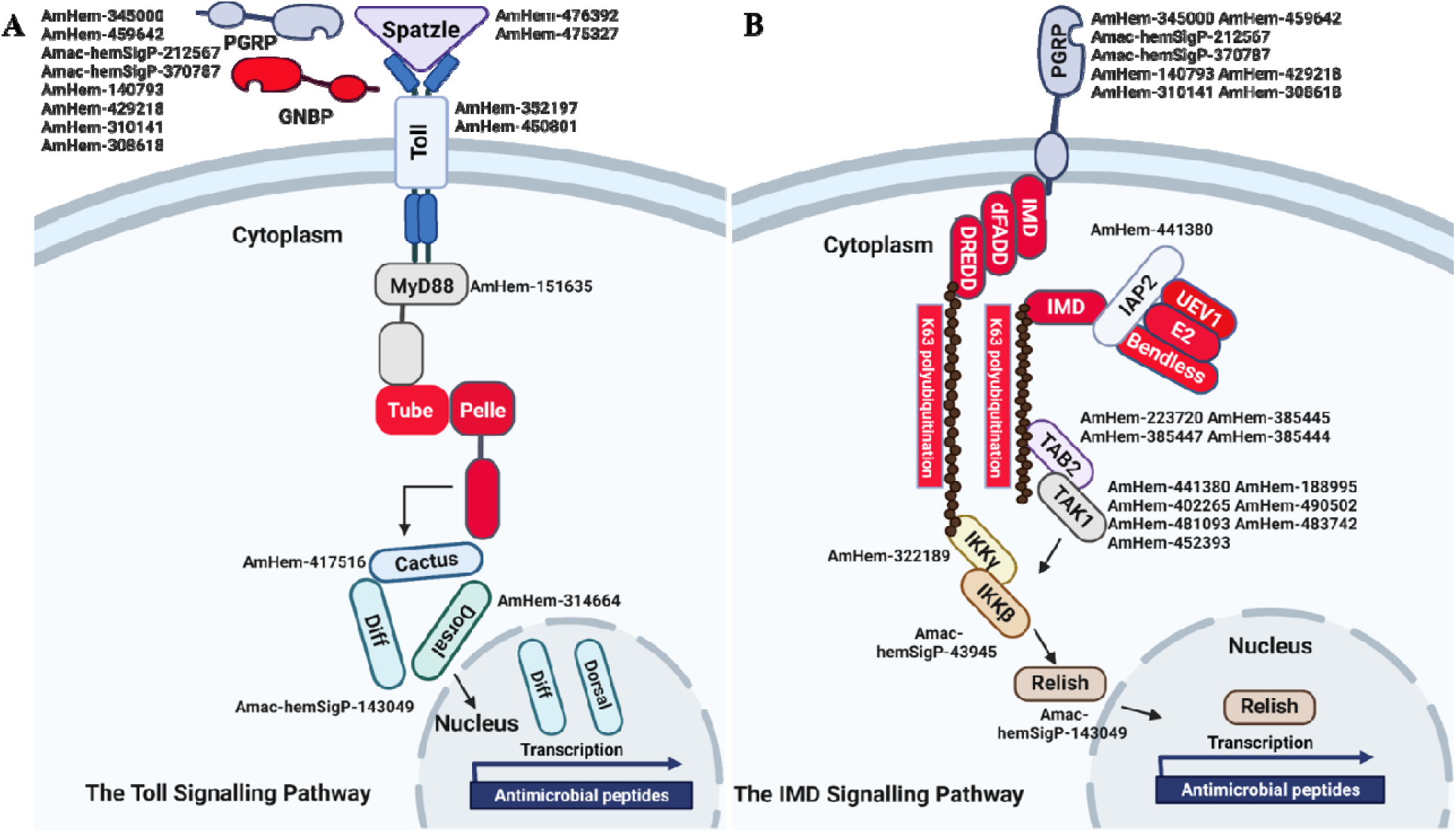
Representative signaling pathways and immune-related genes. Reconstruction of the immune-signaling genes derived from *R. parkeri-*infected and uninfected hemocytes showing the components of the **(A)** Toll and **(B)** IMD signaling pathways. The Toll and IMD signaling pathways are highly conserved in ticks. Transcripts highlighted in red were not identified in this study.

### 3.9 Regulators of the IMD pathway

The immune deficiency (IMD) pathway is activated upon stimulation of PGRPs by the Gram-negative diaminopimelic acid (DAP)-type peptidoglycan, which stimulates both soluble and transmembrane PGRPs. In contrast to the Toll pathway, the IMD pathway contributes to the production of most AMPs in *Drosophila* (79). In our dataset, we identified the differential regulation of several transcripts in the IMD pathway including inhibitor of apoptosis 2 (*IAP2*), mitogen-activated protein kinase-7 (*MAPK7/TAK1*), mitogen-activated protein kinase 7-interacting protein 2 (*MAP3K7IP2/TAB2*), inhibitor of nuclear factor kappa-B kinase subunits (*IKK*), and *Relish*. AmHem-441380 (2-fold upregulated in *R. parkeri-*infected hemocytes) is an *IAP-2* transcript with baculovirus inhibitor of apoptosis protein repeat (*BIR*), ring finger, and zinc finger domains characteristic of IAP proteins. These proteins regulate NF-κB signaling pathways in the cytoplasm (80). MAPK7/TAK1, TAB-2/MAP3K7IP2, and the IKK complex induce cleavage of Relish and subsequent nuclear translocation by transferring a phosphate group to Relish. We identified seven transcripts of *MAPK7/TAK1* (AmHem-188995, AmHem-402265, AmHem-490502, AmHem-481093, AmHem-483742, AmHem-452393, and AmHem-428323) and four *TAB-2/MAP3K7IP2* transcripts (AmHem-223720, AmHem-385445, AmHem-385447, and AmHem-385444), each with STKc and TyrKc domains, which are the catalytic domain of the serine/threonine kinase and tyrosine kinase catalytic domains, respectively (**Fig. 6B, Supplementary Fig. S5**). Infection with *R. parkeri* upregulated of all the *MAPK7/TAK1* transcripts except for AmHem-452393 (12-fold downregulated). Sequences encoding *IMD*, Fas-associated via death domain (*FADD*), and death-related ced-3/Nedd2-like caspase (*DREDD*) genes were absent in our dataset.

### 3.10 Nimrod B2 and eater mediate hemocyte phagocytosis

Hemocytes participate in humoral and cellular defenses in response to microbial infections. Their specific roles are defined by their expressed cell surface receptors. *Nimrod B2* and *eater*, which mediate microbial phagocytosis upon infection (81,82), were downregulated in hemocytes from *R. parkeri-*infected ticks. Their role in hemocyte phagocytosis and as markers of phagocytic hemocytes have been described in hematophagous and non-hematophagous organisms (81–84). We therefore further defined the role of *nimrod B2* (AmHem-305744) and *eater* (AmHem-270031) in hemocyte phagocytosis using a combination of RNAi and *in vivo* phagocytosis approaches (**Fig. 7A**). Quantitative PCR validation of their expression supported the RNA-seq result (**Fig. 7B**). Their transcripts were significantly depleted in dsRNA-injected tick hemocytes (**Fig. 7C**), and the *in vivo* bead phagocytosis assay also revealed depletion in hemocyte phagocytosis of yellow-green FluoSpheres (beads) (**Fig. 7D**). *Nimrod* B2 and *eater* were both significantly downregulated in CLD-depleted hemocytes in our phagocytosis assay (**Fig. 7E and F**), confirming their specificity to phagocytic hemocytes. Together, these data demonstrate that *nimrod B2* and *eater* are two potential candidate marker genes regulating hemocyte phagocytosis.

**Figure 7.**
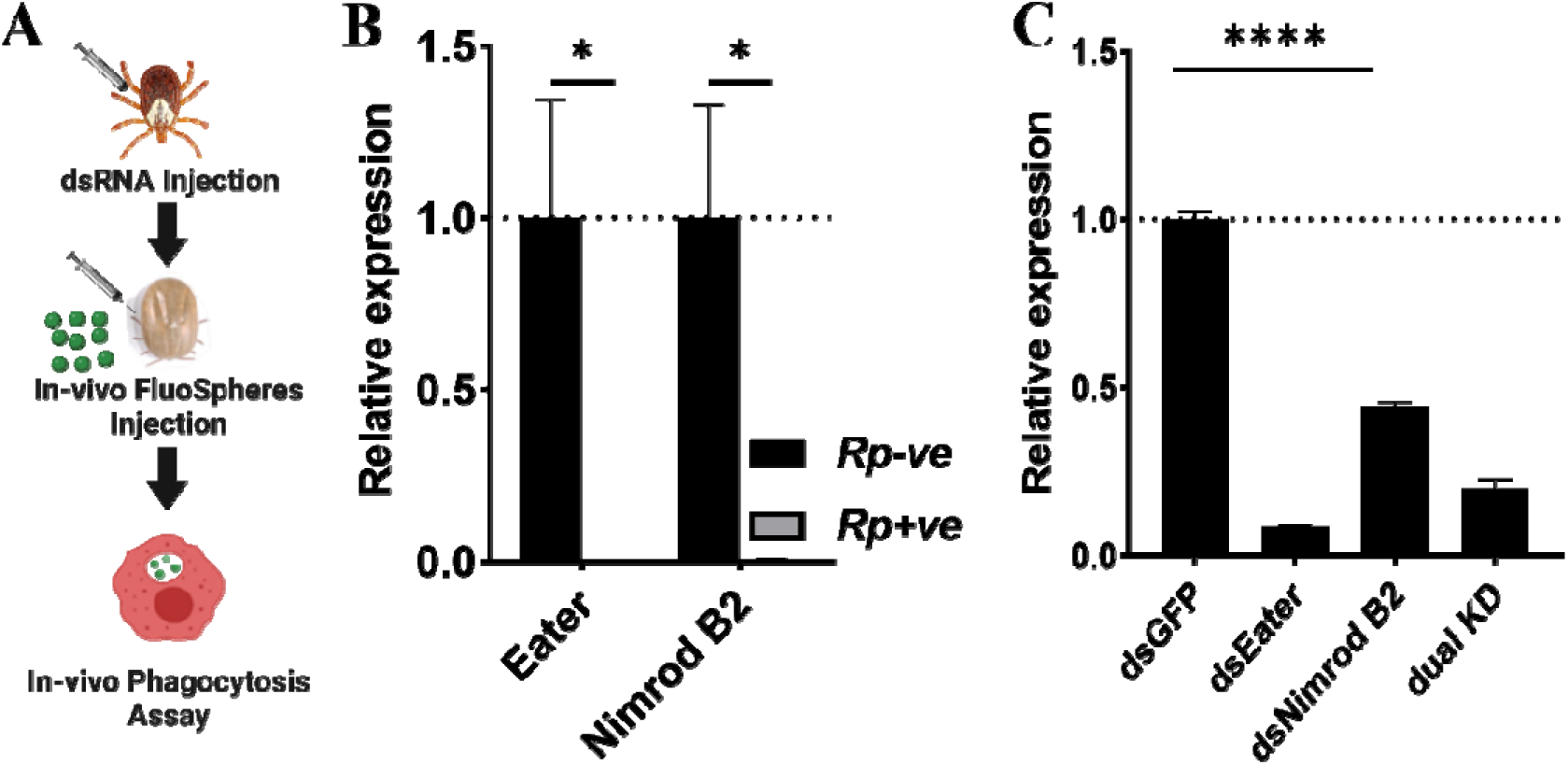
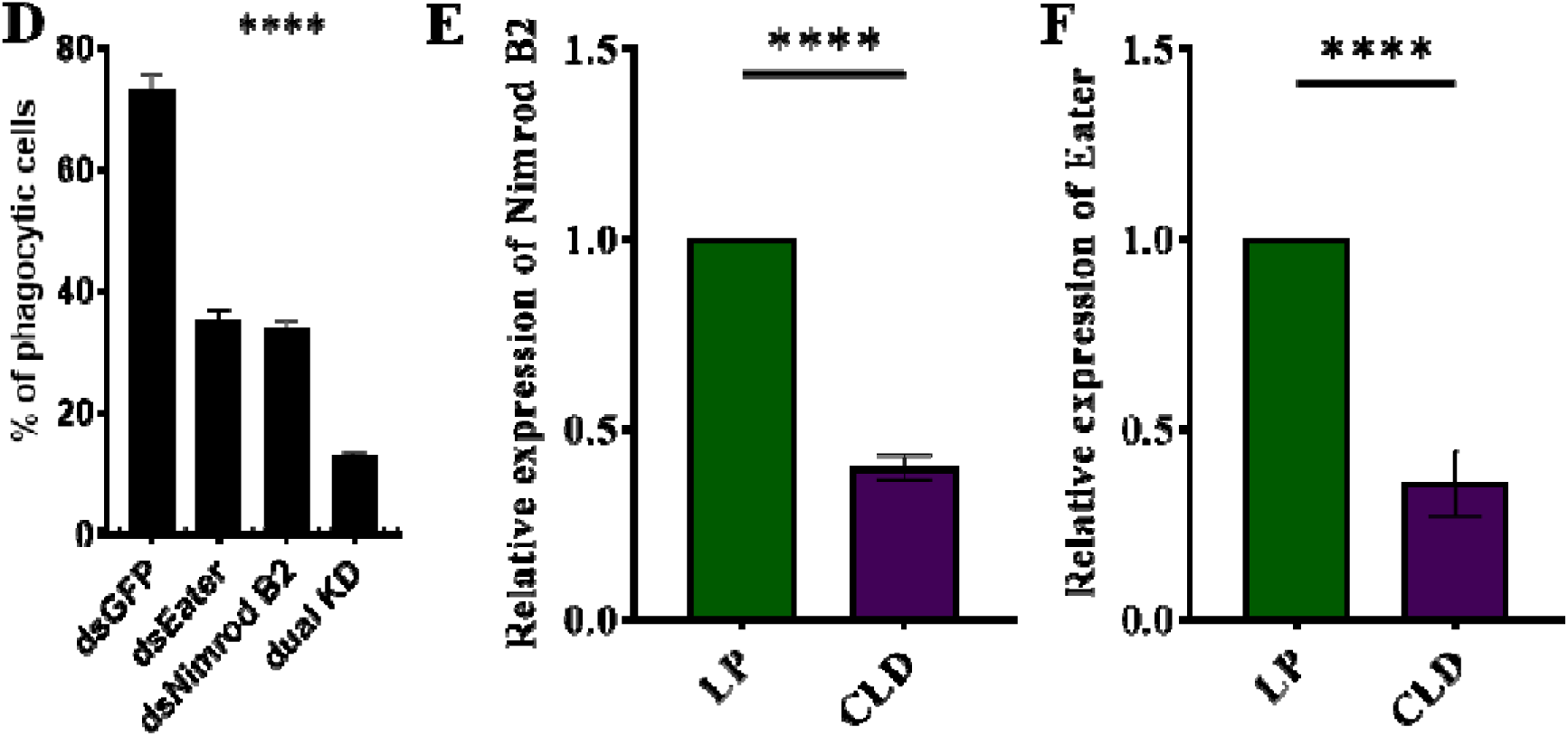
*Nimrod B2* and *eater* as functional markers of hemocyte phagocytosis. The role of *nimrod* B2 and *eater* silencing on *in vivo* phagocytosis was evaluated in *Am. maculatum* hemocytes **(A)**. qPCR validation of bulk RNA expression profiles of *nimrod B2* and *eater* in uninfected and *R. parkeri-*infected *Am. maculatum* hemocytes **(B)**. dsRNA was injected into *Am. maculatum* female ticks to disrupt the expression of *nimrod B2* and *eater* genes and confirmed by qPCR **(C)**. The proportion of phagocytic hemocytes was compared with dsGFP-injected ticks **(D)**. Additional validation of phagocyte depletion showing a significant reduction in *nimrod B2* **(E)** and *eater* **(F)** transcript in CLD-injected ticks. Gene expression was normalized to *Am. maculatum actin*. Data were analyzed using unpaired t-tests in GraphPad Prism v8.4.1. *P < 0.05, **P < 0.01, ***P < 0.001, ****P < 0.0001.

## 4. Discussion

Here we report morphological and functional heterogeneity in *Am. maculatum* hemocytes. We define a role for phagocytic hemocytes in the immune response against bacterial infections and identify potential molecular markers of hemocyte phagocytosis. We report for the first time direct evidence of *R. parkeri* infection of phagocytic hemocytes, which might play a role in the dissemination of these organisms throughout the tick body.

Previous studies have identified different hemocyte subtypes in hard (20,30,31) and soft (3,60) tick species, which vary depending on the developmental stage, infection status, and sex. Here we identified five unique hemocyte subtypes based on histomorphological analysis. Female ticks pose the most threat to human and animal hosts through hematophagy and pathogen transmission, but we did not detect significant differences between hemocyte populations in unfed male and female ticks. Nevertheless, there were functional differences between male and female ticks, with hemocytes from female ticks displaying more phagocytosis than those from males. The functional distinction between male and female ticks could be attributed to the presence of more phagocytic plasmatocytes in female than male ticks. More phagocytic hemocytes may also be needed in female hemolymph due to their large body size relative to males, with the prolonged feeding time on the host of females increasing the chance of microbial growth within the tick, thus necessitating a more robust and primed immune system. Immune priming in invertebrates mimics the vertebrate adaptive immune response upon pathogen infection. While immune system priming is well described in mosquito vectors (85–87), only one study has described the presence of active immune priming in ticks (88). We also observed that blood feeding increased granulocytes in female and plasmatocytes in male ticks.

Although granulocytes are professional phagocytic hemocytes, plasmatocytes have also been shown to display phagocytic functions. Blood feeding also eliminated prohemocytes in both male and female ticks. Prohemocytes are immature hemocytes that can differentiate into other hemocyte types, and their absence would indicate that new hemocytes are produced during feeding, with prohemocytes the source of those new hemocytes. Together, these data for the first time show the heterogeneous nature of the *Am. maculatum* hemocytes and functional differences in male and female hemocytes. Developing specific functional or molecular markers will now be important to confirm our morphological classification.

Chemical inhibitors have been widely used to characterize mammalian immune cells. In the absence of molecular markers to characterize tick hemocytes, hemocytes can be studied and characterized by inhibiting their functions, similar to the widely used reverse genetic approach for gene characterizations. Clodronate liposomes (CLD) are widely used to deplete phagocytic macrophages in mammalian systems (54,55,89). CLD mediates phagocyte killing by releasing toxic clodronate upon macrophage phagocytosis, which subsequently induces apoptosis (54). Phagocytic hemocytes have been depleted with CLD in mosquitoes and *Drosophila* (32-33). We also showed that CLD can successfully reduce the phagocytic hemocyte population in *Am. maculatum*, as evidence by a reduced proportion of phagocytic hemocytes and subsequent loss in functional phagocytosis confirmed by immunofluorescence on CLD treatment. These findings were further supported by a decrease in *nimrod B2* and *eater* transcript in CLD-treated ticks, both of which are well characterized phagocyte markers in mosquitoes (33, 83) and *Drosophila* (81-82, 84). In the tick system, we showed that *nimrod B2* and *eater* knockdown significantly reduced the proportion of phagocytic hemocytes, further suggesting that these two genes may be useful candidate markers of phagocytic hemocytes in *Am. maculatum*. By showing successful depletion of phagocytic hemocytes using CLD, these experiments provide the means to functionally characterize phagocytic hemocytes in *Am. maculatum* and other tick species. Our experiments also serve as proof of concept for using CLD in the functional study of phagocytic hemocytes in a non-model organism such as ticks. Due to the lack of specific antibodies targeting these two genes, further characterization of their functions is limited.

As the first component of the cellular immune response, phagocytic hemocytes are primarily responsible for scavenging and removing invading pathogens, mainly via phagocytosis. The reduced survival of CLD-depleted ticks following bacterial challenge further highlights the critical role of hemocytes in maintaining immune responses in ticks, consistent with immune impairments following CLD depletion seen in mosquitoes and *Drosophila* (32-33). While ours is the first functional validation of phagocytic hemocytes by CLD depletion in ticks, two studies used alternative approaches to inhibit hemocyte function and demonstrated reduced survival in ticks following bacterial challenge (6, 11). The outcome of our experiments and similar reports in other organisms highlight the functional conservation of these immune cells in invertebrates.

Hemolymph is a well-defended niche containing hemocytes and several soluble effector molecules that directly inactivate or kill invading microbes (90,91). Pathogens acquired via an infected host’s blood must leave the blood bolus and infect the midgut and resident tissues. The process of *R. parkeri* dissemination from the point of midgut infection to other tick tissues is not fully understood. It has been proposed that rickettsiae can cross the midgut barrier to infect hemocytes during blood feeding (92), and others have also demonstrated *Rickettsia* infection of the tracheal system (93). The constant bathing of tissues with hemolymph and the persistence of the tracheal system through developmental stages make them viable routes for the systemic maintenance and dissemination of rickettsiae throughout the tick body. In the current study, we detected rod-shaped *R. parkeri* in the hemocytes of naturally infected *Am. maculatum*. Likewise, capillary feeding and microinjection of GFP-expressing *R. parkeri* also led to the observation of *Rickettsia* organisms in hemocytes, suggesting active infection of hemocytes by the pathogen. Dissemination of *R. parkeri* to the *Am. maculatum* midgut, salivary gland, and ovarian tissues has previously been described following capillary feeding (94); however, dissemination into the hemolymph and infection of hemocytes is a novel observation that argues for a role for circulating hemocytes in rickettsial trafficking throughout the tick body. Our detection of *R. parkeri* in the hemocytes of infected ticks and tissue dissemination of *R. parkeri* in *Am. maculatum* salivary glands, midguts, and ovaries reported by Harris et al. (94) and (95) strongly suggest that the pathogen survives long enough in the hostile hemolymph environment to find its way to other tick organs. *R. parkeri* and other vector-borne diseases have evolved a complex process enabling them to colonize and disseminate throughout the arthropod host through transovarial and transstadial transmission (39, 40, 96, 97). Similar to this, salivary gland colonization during feeding is a crucial step for infecting the mammalian host (98-99). The presence of sessile hemocytes (tissue-associated hemocytes) has been described in mosquitoes and *Drosophila* (100-101), and these hemocytes are found in those regions of the body that interact most with invading pathogens such as the periosteal region and abdominal walls (100). However, their role in transovarial and transstadial maintenance of pathogens remains unknown. Whether hemocytes play a role in the trafficking of pathogenic bacteria to tick tissues as observed for mosquito phagocytes and viral infection still needs to be established (61). The mechanisms by which *R. parkeri* enter and survive inside hemocytes also require further examination.

Molecular studies of tick hemocyte biology and hemocyte-mediated immune responses have been limited due to, in part, the technical challenges surrounding hemolymph collection from different tick stages, the lack of a hemocyte-like cell line (as with *Drosophila* and mosquitoes), and a lack of hemocyte-specific markers. To identify the molecular responses in *Am. maculatum* hemolymph during infection, hemocyte transcriptomes from *R. parkeri-*infected and uninfected ticks were isolated and analyzed. We identified an entire repertoire of transcripts differentially expressed in hemocytes on *R. parkeri* infection (**Supplementary Table S2**). We identified several genes that mediate the cellular immune response, especially genes encoding hematopoietic functions and hemocyte differentiation. The anatomical structure and molecular basis of hemocyte production in ticks are still unknown. An earlier study on hemolymph circulation proposed the presence of a lymph-like organ as the site of hemolymph production in ticks (14), but the active production of hemocytes from the tick lymph gland or a specialized hematopoietic organ has yet to be demonstrated. The presence of transcriptional and humoral regulators of hematopoiesis genes suggests that the regulation of hematopoiesis is conserved in arthropod and insect species. For instance, the transcription factors *GATA* and *Runt* were detected in our data, which are critical for the proliferation of hematopoietic stem cells and differentiation of myeloid stem cells, respectively, in invertebrates (102). *Lozenge*, a Runt homolog, has been reported to mediate crystal cell maturation in *Drosophila* (103, 104), while in crayfish it mediates differentiation from hematopoietic stem cells to mature granular and semigranular cells (64). Kwon et al. also demonstrated a reduction in oenocytoid differentiation following silencing of *Lozenge* in *An. gambiae* mosquitoes (56) as well as a decrease in the expression of *PPO3/8*, which are molecular markers of oenocytoid cells (56). *Serpent* is a *Drosophila* GATA factor that regulates hematopoiesis and is an ortholog of the vertebrate GATA family (105). *Serpent* is expressed in hemocyte precursors but, unlike *lozenge*, it is still expressed in mature prohemocytes and crystal cells, suggesting roles beyond hematopoiesis (106). No studies have yet identified *Runt* or its homolog in any tick species; however, a GATA factor has been described in *Haemaphysalis longicornis* that activates *Vitellogenin*, which is essential for reproduction (107–109). The presence of transcripts that encode humoral regulators of hematopoiesis, such as *Astakines*, β*-thymosin*, and *transglutaminases*, was an unexpected finding. Astakines regulate hematopoiesis via interaction with transglutaminases or via direct interactions with hematopoietic cells to promote structural rearrangements (64). Proteins containing thymosin domains have been described in *Drosophila*, where Ciboulot, a protein with three thymosin domains, regulates axon growth during brain metamorphosis (110). Thypedin, a protein with 27 thymosin domains, regulates foot regeneration in Hydra (111). Ovarian expression of β*-thymosin* was reported in the dog tick, *Dermacentor variabilis* (112). In invertebrates, TGases were initially described as coagulation factors and later shown to regulate hematopoiesis in crustaceans (113, 114) by maintaining hematopoietic cells in an undifferentiated state, thus suppressing hematopoiesis (115, 116). It is currently unclear how these genes regulate the production and differentiation of hemocytes in ticks. However, the presence of some of these genes, such as *Astakines*, in the genomes of crustaceans, ticks, scorpions, spiders, and other arthropods but not *Drosophila* or mosquitoes suggests evolutionary divergence in hematopoietic processes (117).

Several PGRP-encoding transcripts were detected, consistent with a previous hemocyte transcriptome study in *Ixodes scapularis*, where twenty-six PGRP-encoding sequences were detected (62). Genes encoding the Toll and IMD pathways have previously been reported in ticks, although some components of these pathways have yet to be identified (88, 118–122). While many genes encoding Toll and IMD pathway components were identified in our data, it is unsurprising that some Toll and IMD pathway components were not identified in this study. For instance, the absence of *GNBP, Tube*, and *Pelle* transcripts-components of the Toll pathway - and the IMD components *IMD, FADD*, and *DREDD* is supported by the absence of these transcripts from the *Ix. scapularis* and *Rhipicephalus microplus* genomes (118, 123). Shaw and colleagues described plasticity in the immune pathways between insects and arthropods, demonstrating activation of the *Ix. scapularis* IMD pathway by two infection-derived glycerol following pathogen infection (88). Our knowledge of tick immunity has hitherto relied on model organisms like *Drosophila* and mosquitoes, but our data and that from recent studies on tick immunity argue that immunity in hematophagous arthropods such as ticks is very different (88, 118, 123).

Cellular immunity relies on hemocytes directly killing invading microbes via phagocytosis, melanization, or the production of reactive oxygen species. Although there have been many studies of cellular immunity in ticks, the roles of specific immune cell types, their molecular signatures, and their contribution to the hemocyte-mediated immune response are still not fully understood. Our study identifies a repertoire of potential candidate genes that regulate hemocyte functions. An unexpected observation was the identification of *nimrod B2* and *eater* transcripts, which were significantly expressed in uninfected ticks but downregulated on *R. parkeri* infection. Nimrod B2 and eater are both transmembrane receptors expressed on hemocyte membranes shown to be responsible for the phagocytosis of Gram-positive and Gram-negative bacteria in *Drosophila* (81-82, 84) and mosquitoes (33, 83, 124). Although the precise roles of nimrod B2 and eater in tick immunity are unknown, our RNAi experiments suggest that they play a direct role in hemocyte phagocytosis. Further studies are now required to determine the cellular localization of these two genes and whether their expression is unique to phagocytic hemocytes from other tick species. Our study also confirms the presence of an active complement component system with several TEPs. The tick complement system is essential for killing tick-transmitted pathogens, as demonstrated by Urbanová and colleagues (4), who demonstrated phagocytosis of the spirochete *Borrelia afzelli* in *Ix. ricinus*. A similar role for tick complement-like proteins in the phagocytosis of non-pathogenic microbes was also demonstrated in *Ix. ricinus* ticks (4,125,126).

## 5. Conclusion

Here we describe morphological and functional differences in *Am. maculatum* hemocytes and characterize transcriptional changes in cellular and humoral responses to *R. parkeri* infection. Our results reveal heterogenous hemocyte populations in *Am. maculatum* showing variable phagocytic capacity. We also describe an integral role for phagocytes in responses to microbial pathogens. We for the first time observed *R. parkeri* in phagocytic hemocytes. Our transcriptome analysis of *R. parkeri-*infected and uninfected hemocytes allowed us to explore differentially expressed immunity and hematopoietic genes on *R. parkeri* infection. This “big” transcriptome dataset will be important for identifying potential biomarkers of hemocyte subtype, function, and production. Our findings also raise important questions about the role of each hemocyte subtype in immune responses and vector competence.

## 6. Data Availability Statement

The raw fastq files were deposited in the Sequence Read Archives (SRA) of the National Center Biotechnology Information (NCBI) under bioproject PRJNA878782 and biosamples SAMN30755417 and SAMN30755418. Deduced coding sequences and their translations were deposited to the Transcriptome Shotgun Assembly database DDBJ/EMBL/GenBank under the accessions GKCB01000001-GKCB01011171.

## 7. Ethics Statement

Protocols for tick blood-feeding were approved by the University of Southern Mississippi’s Institutional Animal Care and Use Committee (USMIACUC protocols #15101501.3 and 17101206.2).

## SUPPLEMENTARY DATA

**Supplementary Figure S1A-C.**
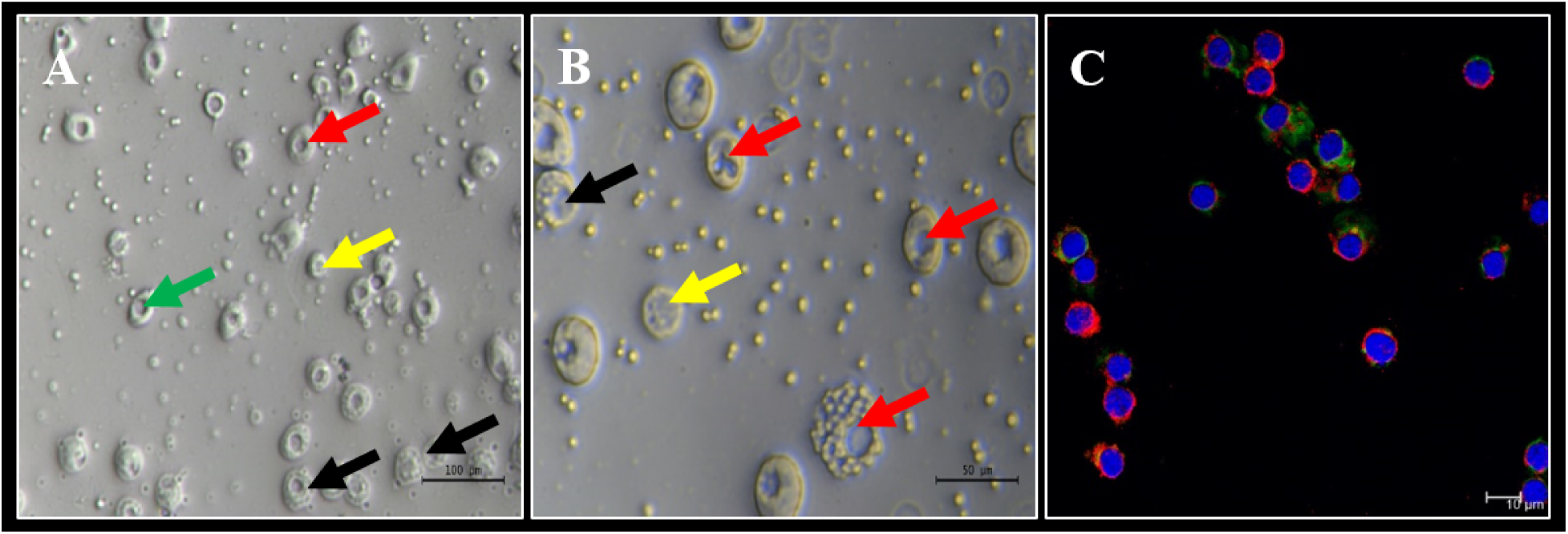
Microscopic examination and immunostaining of hemolymph. Light microscopic examination of perfused hemolymph at **(A)** lower and **(B)** higher magnification showing multiple hemocyte populations. **(C)** Immunostaining of perfused hemocytes with WGA lectin (green), Vybrant CM-Dil (red), and Hoechst 33342 stains (blue). Scale bars as indicated.

**Supplementary Figure S1D-I.**
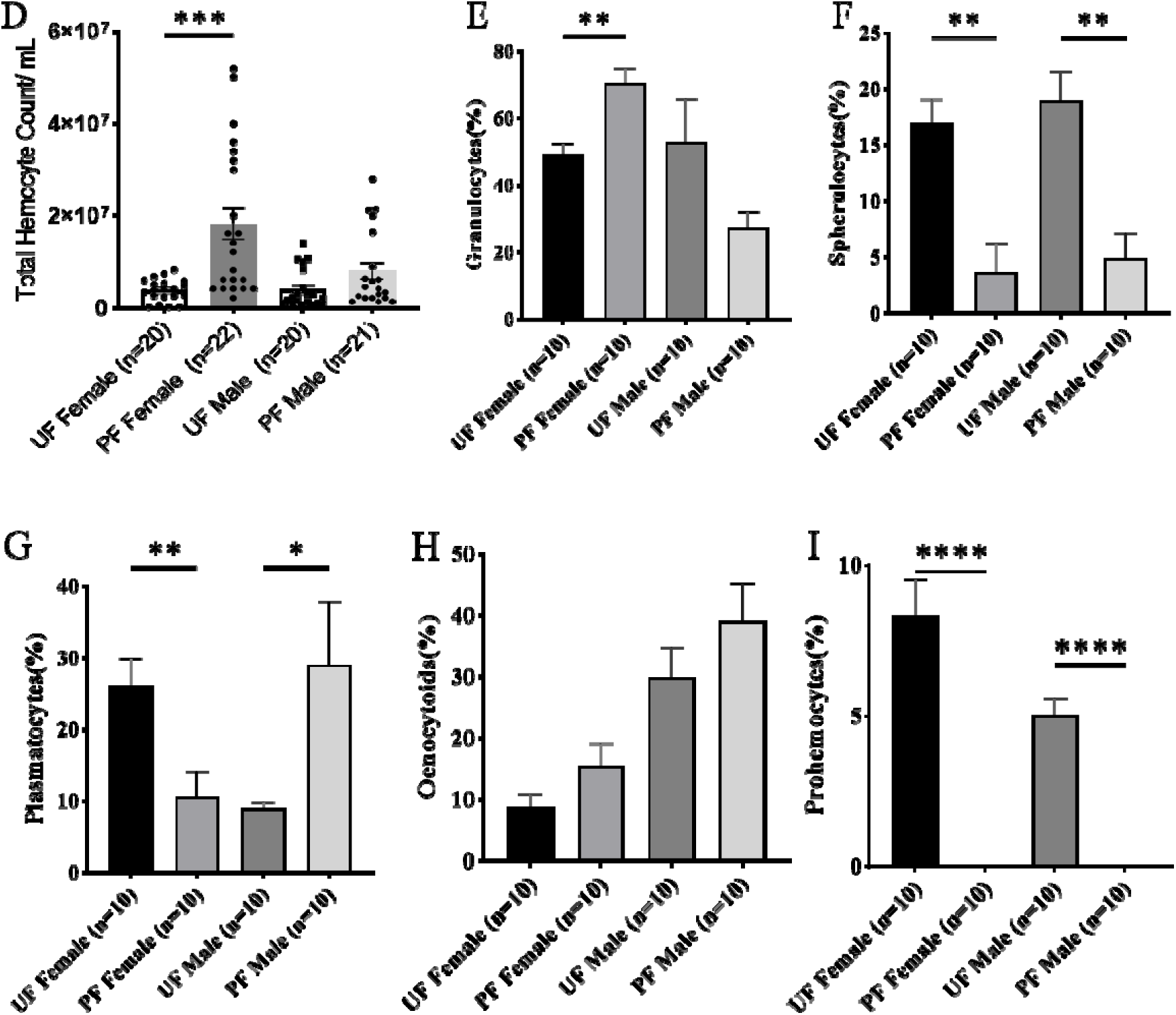
Hemocyte populations in unfed ticks. Hemolymph was perfused from unfed male and female ticks and the **(D)** total and **(E-I)** differential hemocyte populations compared between unfed and partially fed male and female ticks using an improved Neaubeur chamber. Data were analyzed using unpaired t-tests in GraphPad Prism v8.4.1. *P < 0.05, **P < 0.01, ***P < 0.001, ****P < 0.0001. UF; unfed, PF; partially blood fed.

**Supplementary Figure S2A and B.**
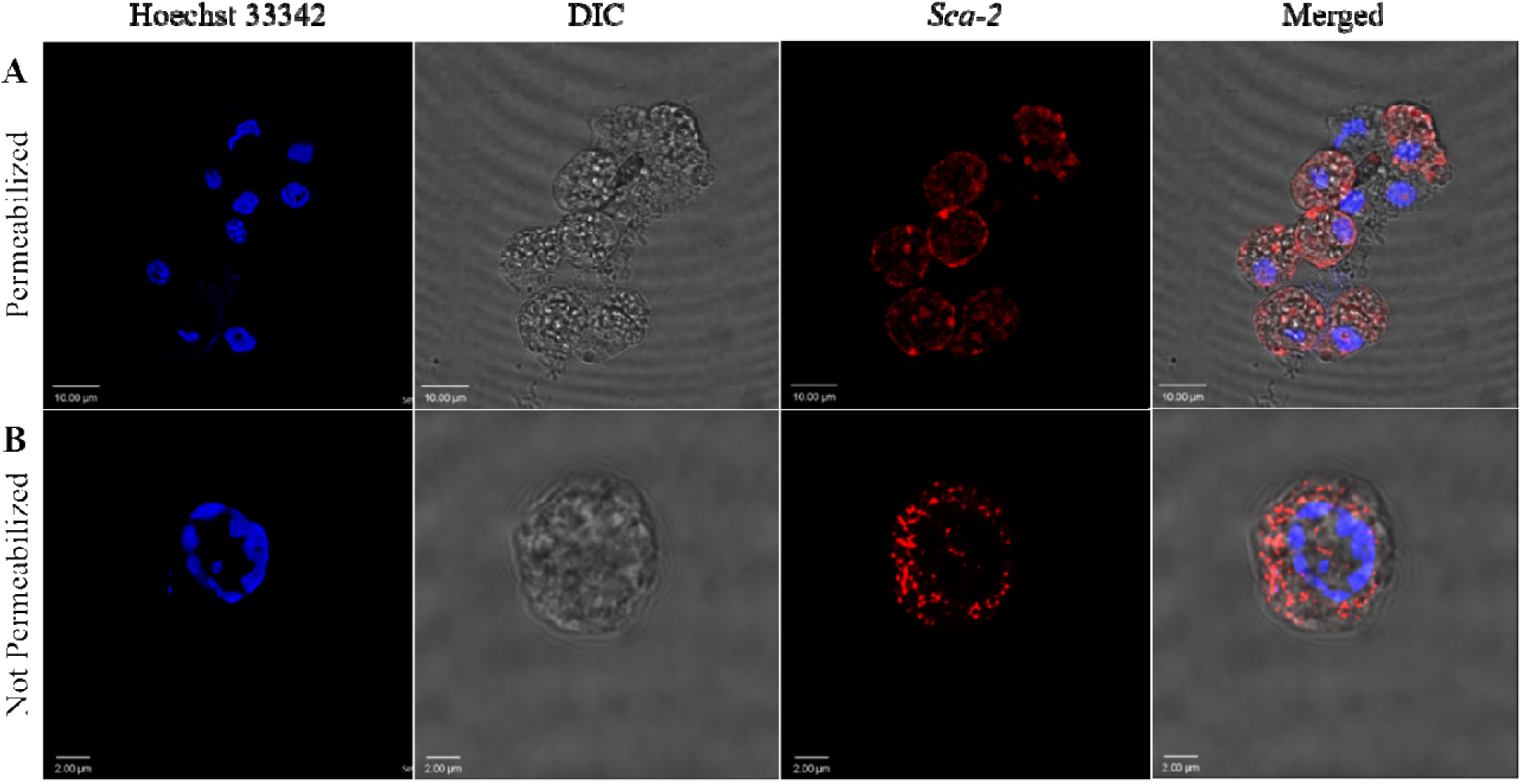
Confocal images of phagocytic hemocytes infected with *Rickettsia parkeri*. Representative confocal images of immunofluorescence staining for *R. parkeri Sca-2* protein in hemolymph of *R. parkeri-*infected *Am. maculatum*. Hemocytes were either **(A)** permeabilized with Triton-X or **(B)** not permeabilized before incubation with *Sca-2-* specific antibodies (red) and Hoechst 33342 (blue).

**Supplementary Figure S3A and B.**
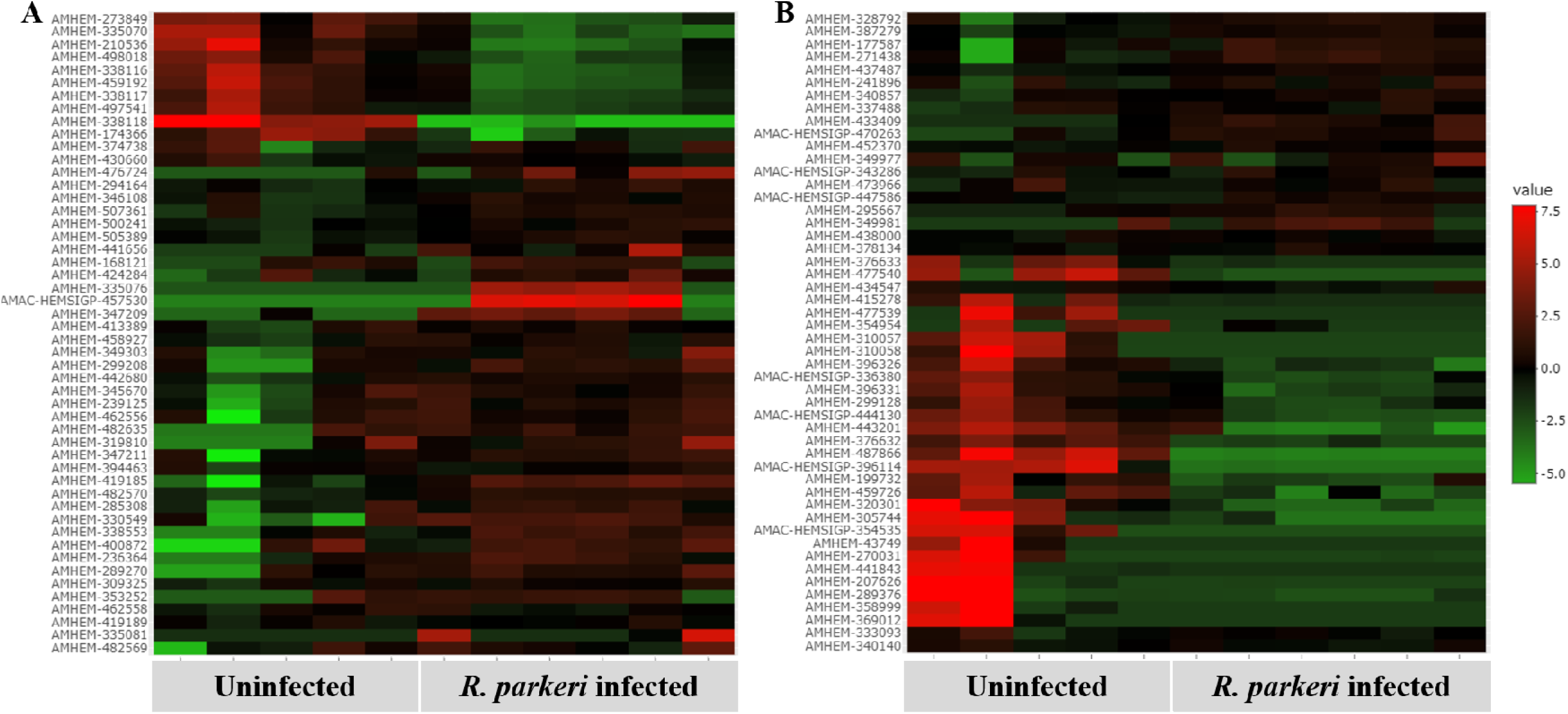
Heatmaps of RNA-seq expression data showing hematopoietic and cellular function genes differentially regulated in *R. parkeri-*infected compared with uninfected hemocytes. Several of these transcripts include **(A)** transcriptional and **(B)** humoral regulators of hemocyte differentiation and maturation. The scale bar shows the log_2_fold expression value of the transcripts.

**Supplementary Figure S4.**
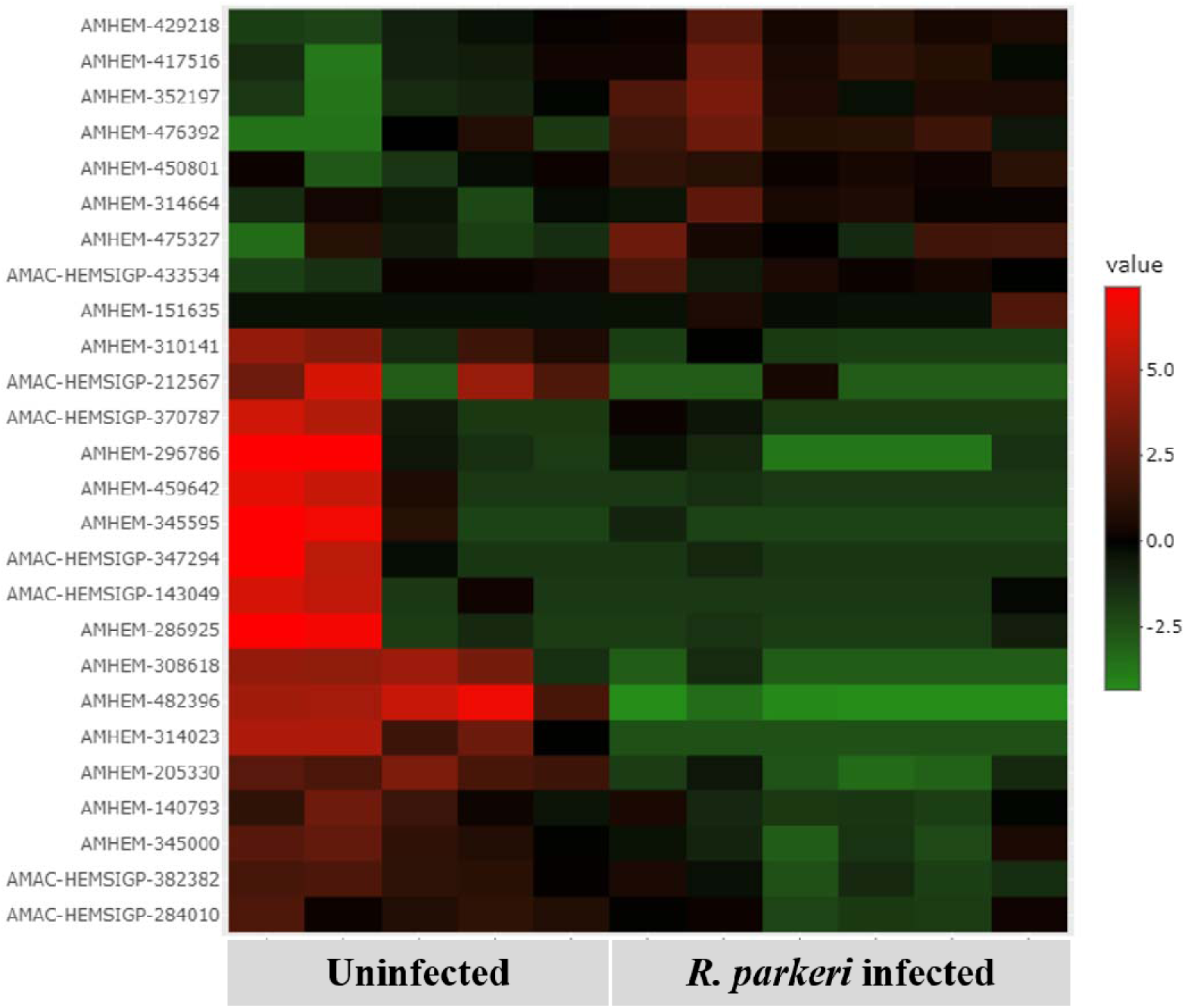
Heatmap of RNA-seq expression data showing differentially expressed transcripts in the toll-signaling pathway in *R. parkeri* infected and uninfected hemocytes. Six of the nine Toll genes were identified including eight *PGRP* transcripts. Transcripts encoding *GNBP, tube*, and *pelle* were, however, not present. The scale bar to the right shows the log_2_fold expression value of the transcripts.

**Supplementary Figure S5.**
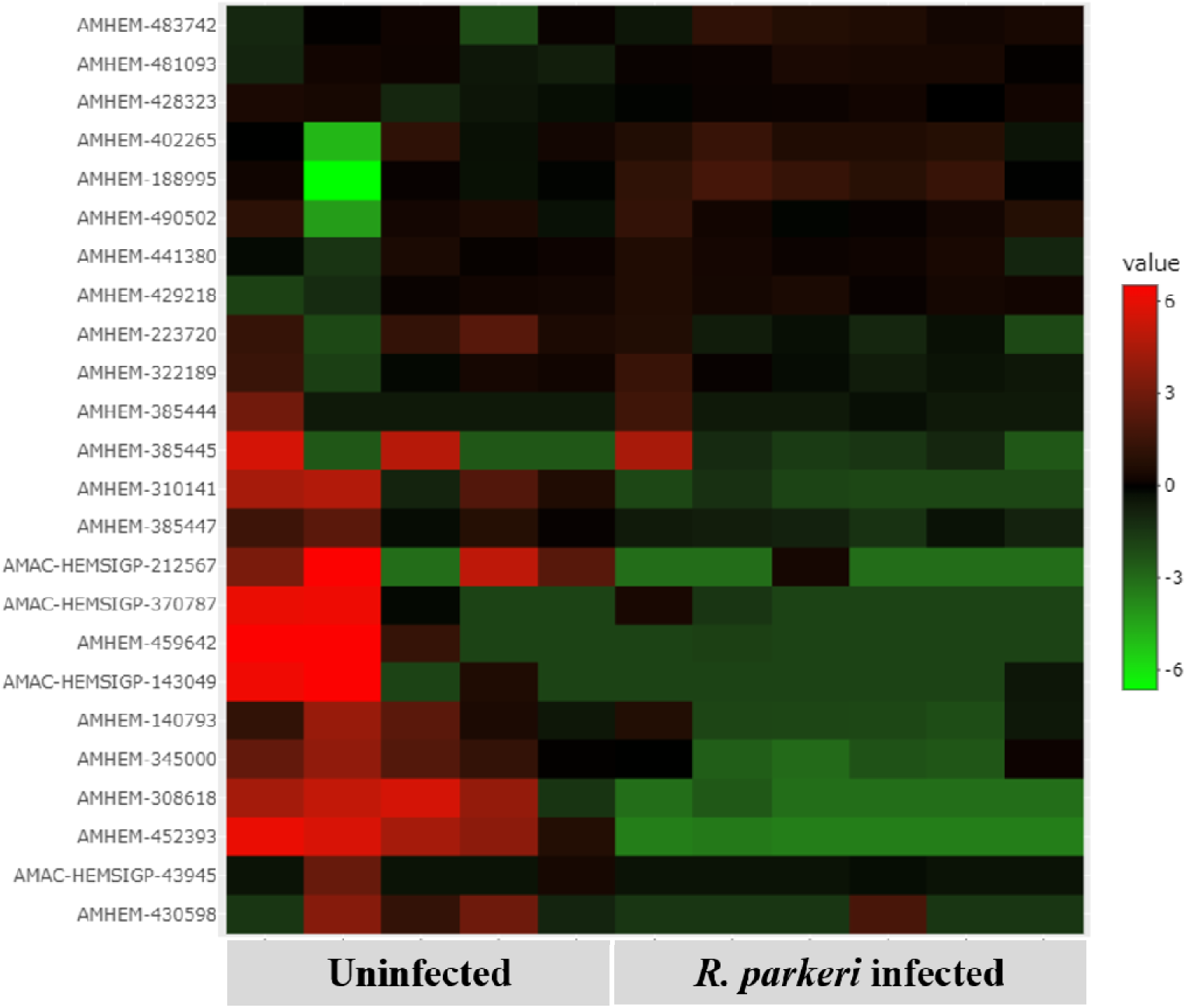
Heatmap of RNA-seq expression data showing differentially expressed transcripts in the IMD signaling pathway in *R. parkeri-*infected and uninfected hemocytes. Sequences encoding *IMD*, Fas-associated via death domain (*FADD*). and deathrelated ced-3/Nedd2-like caspase (*DREDD*) genes were absent from our data. The scale bar to the right shows the log_2_fold expression value of the transcripts.

## Notes

### Competing Interest Statement

The authors have declared no competing interest.

## References

1. Biggs HM, Behravesh CB, Bradley KK, Dahlgren FS, Drexler NA, Dumler JS, et al. Diagnosis and Management of Tickborne Rickettsial Diseases: Rocky Mountain Spotted Fever and Other Spotted Fever Group Rickettsioses, Ehrlichioses, and Anaplasmosis - United States. MMWR Recomm reports Morb Mortal Wkly report Recomm reports. 2016;65(2):1–44.

2. Sumner JW, Durden LA, Goddard J, Stromdahl EY, Clark KL, Reeves WK, et al. Gulf Coast Ticks (Amblyomma maculatum) and Rickettsia parkeri, United States. Emerg Infect Dis. 2007;13(5):751.

3. Inoue N, Hanada K, Tsuji N, Igarashi I, Nagasawa H, Mikami T, et al. Characterization of Phagocytic Hemocytes in Ornithodoros moubata (Acari: Ixodidae). J Med Entomol. 2001 Jul 1;38(4):514–9.

4. Urbanová V, Hajdušek O, Mondeková HH, Šíma R, Kopáček P. Tick thioester-containing proteins and phagocytosis do not affect transmission of Borrelia afzelii from the competent vector Ixodes ricinus. Front Cell Infect Microbiol. 2017 Mar 16;7(MAR):73.

5. Mondekova HH, Sima R, Urbanova V, Kovar V, Ryan RO, Grubhoffer L, et al. Characterization of Ixodes ricinus fibrinogen-related proteins (Ixoderins) discloses their function in the tick innate immunity. Front Cell Infect Microbiol. 2017 Dec 8;7(DEC):509.

6. Aung KM, Boldbaatar D, Umemiya-Shirafuji R, Liao M, Tsuji N, Xuenan X, et al. Hlsrb, a class b scavenger receptor, is key to the granulocyte-mediated microbial phagocytosis in ticks. PLoS One. 2012 Mar 29;7(3).

7. Feitosa APS, Chaves MM, Veras DL, de Deus DMV, Portela NC, Araújo AR, et al. Assessing the cellular and humoral immune response in Rhipicephalus sanguineus sensu lato (Acari: Ixodidae) infected with Leishmania infantum (Nicolle, 1908). Ticks Tick Borne Dis. 2018 Sep 1;9(6):1421–30.

8. Pereira LS, Oliveira PL, Barja-Fidalgo C, Daffre S. Production of reactive oxygen species by hemocytes from the cattle tick Boophilus microplus. Exp Parasitol. 2001;99(2):66–72.

9. Fogaça AC, Almeida IC, Eberlin MN, Tanaka AS, Bulet P, Daffre S. Ixodidin, a novel antimicrobial peptide from the hemocytes of the cattle tick Boophilus microplus with inhibitory activity against serine proteinases. Peptides. 2006 Apr;27(4):667–74.

10. Kocan KM, De La Fuente J, Manzano-Roman R, Naranjo V, Hynes WL, Sonenshine DE. Silencing expression of the defensin, varisin, in male Dermacentor variabilis by RNA interference results in reduced Anaplasma marginale infections. Exp Appl Acarol. 2008 Dec;46(1–4):17–28.

11. Fiorotti J, Urbanová V, Gôlo PS, Bittencourt Vrep, Kopáček P. The role of complement in the tick cellular immune defense against the entomopathogenic fungus Metarhizium robertsii. Dev Comp Immunol. 2022 Jan 1;126:104234.

12. Fogaça AC, Almeida IC, Eberlin MN, Tanaka AS, Bulet P, Daffre S. Ixodidin, a novel antimicrobial peptide from the hemocytes of the cattle tick Boophilus microplus with inhibitory activity against serine proteinases. Peptides. 2006 Apr 1;27(4):667–74.

13. Rego ROM, Kovář V, Kopáček P, Weise C, Man P, Šauman I, et al. The tick plasma lectin, Dorin M, is a fibrinogen-related molecule. Insect Biochem Mol Biol. 2006 Apr 1;36(4):291–9.

14. Binnington KC, Obenchain FD. Structure and Function of the Circulatory, Nervous, and Neuroendocrine Systems of Ticks. In: Physiology of Ticks. Elsevier; 1982. p. 351–98.

15. Biology of Ticks Volume 1 - Daniel E. Sonenshine; R. Michael Roe - Oxford University Press [Internet]. [cited 2020 Sep 2]. Available from: https://global.oup.com/academic/product/biology-of-ticks-volume-1-9780199744053?cc=us&lang=en&

16. Borovičková B, Hypša V. Ontogeny of tick hemocytes: A comparative analysis of Ixodes ricinus and Ornithodoros moubata. Exp Appl Acarol. 2005 Apr;35(4):317–33.

17. Kuhn KH, Haug T. Ultrastructural, cytochemical, and immunocytochemical characterization of haemocytes of the hard tick Ixodes ricinus (Acari; Chelicerata). Cell Tissue Res. 1994 Sep;277(3):493–504.

18. Zhioua E, Lebrun RA, Johnson PW, Ginsberg HS. Ultrastructure of the haemocytes of Ixodes scapularis (Acari: Ixodidae). Acarologia. 1996;37(3):173–9.

19. Sandra B da Silva, Graziela Savastano VREPB. Cellular types involved in the immune response of engorged females of Boophilus microplus inoculated with Metarhizium anisopliae and Penicillium sp. Rev Bras Med Veterinária. 2006;15:128–31.

20. Brinton LP, Burgdorfer W. Fine structure of normal hemocytes in Dermacentor andersoni Stiles (Acari:Ixodidae). J Parasitol. 1971;57(5):1110–27.

21. Kuhn KH, Rittig M, Häupl T, Burmester GR. G1.P2 Haemocytes of the hard tick Ixodes ricinus express coiling phagocytosis of Borrelia burgdorferi. In: Developmental and Comparative Immunology. Elsevier Ltd; 1994.

22. Rittig MG, Kuhn KH, Dechant CA, Gauckler A, Modolell M, Ricciardi-Castagnoli P, et al. Phagocytes from both vertebrate and invertebrate species use “coiling” phagocytosis. Dev Comp Immunol. 1996 Nov 1;20(6):393–406.

23. Liu L, Narasimhan S, Dai J, Zhang L, Cheng G, Fikrig E. Ixodes scapularis salivary gland protein P11 facilitates migration of Anaplasma phagocytophilum from the tick gut to salivary glands. EMBO Rep. 2011 Nov;12(11):1196–203.

24. Hillyer JF. Insect immunology and hematopoiesis. Dev Comp Immunol. 2016;

25. Hillyer JF. Mosquito Immunity. Adv Exp Med Biol. 2010;708:218–38.

26. Nakhleh J, El Moussawi L, Osta MA. The Melanization Response in Insect Immunity. Adv In Insect Phys. 2017 Jan 1;52:83–109.

27. Patrick CD, Hair JA. Laboratory rearing procedures and equipment for multi host ticks (Acarina: Ixodidae). J Med Entomol. 1975;12(3):389–90.

28. Patton TG, Dietrich G, Brandt K, Dolan MC, Piesman J, Gilmore RD. Saliva, salivary gland, and hemolymph collection from Ixodes scapularis ticks. J Vis Exp. 2012 Feb 21;60(60):3894.

29. Aguilar-Díaz H, Quiroz-Castañeda RE, Salazar-Morales K, Miranda-Miranda E. A newly optimized protocol to extract high-quality hemolymph from the cattle tick Rhipicephalus microplus: Improving the old conditions. Curr Res Parasitol Vector-Borne Dis. 2022 Jan 1;2:100066.

30. Fiorotti J, Menna-Barreto RFS, Gôlo PS, Coutinho-Rodrigues CJB, Bitencourt ROB, Spadacci-Morena DD, et al. Ultrastructural and Cytotoxic Effects of Metarhizium robertsii Infection on Rhipicephalus microplus Hemocytes. Front Physiol. 2019 May 29;10(MAY):654.

31. Feitosa APS, Alves LC, Chaves MM, Veras DL, Silva EM, Aliança ASS, et al. Hemocytes of Rhipicephalus sanguineus (Acari: Ixodidae): Characterization, Population Abundance, and Ultrastructural Changes Following Challenge with Leishmania infantum. J Med Entomol. 2015 Nov 1;52(6):1193–202.

32. Kumar JR, Smith JP, Kwon H, Smith RC, Dionne MS, Smith RC. Use of Clodronate Liposomes to Deplete Phagocytic Immune Cells in Drosophila melanogaster and Aedes aegypti. Front Cell Dev Biol. 2021 Feb 2;9:627976.

33. Kwon H, Smith RC. Chemical depletion of phagocytic immune cells in Anopheles gambiae reveals dual roles of mosquito hemocytes in anti-Plasmodium immunity. Proc Natl Acad Sci U S A. 2019;116(28):14119–28.

34. Ammerman NC, Beier-Sexton M, Azad AF. Laboratory Maintenance of Rickettsia rickettsii. Curr Protoc Microbiol. 2008 Nov 1;11(1):3A.5.1-3A.5.21.

35. Smith RC, Barillas-Mury C, Jacobs-Lorena M. Hemocyte differentiation mediates the mosquito late-phase immune response against Plasmodium in Anopheles gambiae. Proc Natl Acad Sci U S A. 2015;112(26):E3412–20.

36. Salic A, Mitchison TJ. A chemical method for fast and sensitive detection of DNA synthesis in vivo. Proc Natl Acad Sci U S A. 2008 Feb 19;105(7):2415–20.

37. Bryant WB, Michel K. Blood feeding induces hemocyte proliferation and activation in the African malaria mosquito, Anopheles gambiae Giles. J Exp Biol. 2014 Apr 1;217(8):1238–45.

38. Crispell G, Budachetri K, Karim S. Rickettsia parkeri colonization in Amblyomma maculatum: the role of superoxide dismutases. Parasit Vectors. 2016 Dec 20;9(1):291.

39. Budachetri K, Karim S. An insight into the functional role of thioredoxin reductase, a selenoprotein, in maintaining normal native microbiota in the Gulf Coast tick (Amblyomma maculatum). Insect Mol Biol. 2015 Oct 1;24(5):570–81.

40. Budachetri K, Kumar D, Crispell G, Beck C, Dasch G, Karim S. The tick endosymbiont Candidatus Midichloria mitochondrii and selenoproteins are essential for the growth of Rickettsia parkeri in the Gulf Coast tick vector. Microbiome. 2018 Aug 13;6(1).

41. Kumar D, Budachetri K, Meyers VC, Karim S. Assessment of tick antioxidant responses to exogenous oxidative stressors and insight into the role of catalase in the reproductive fitness of the Gulf Coast tick, Amblyomma maculatum. Insect Mol Biol. 2016 Jun 1;25(3):283.

42. Bullard RL, Williams J, Karim S. Temporal Gene Expression Analysis and RNA Silencing of Single and Multiple Members of Gene Family in the Lone Star Tick Amblyomma americanum. PLoS One. 2016 Feb 1;11(2):e0147966.

43. Simpson JT, Wong K, Jackman SD, Schein JE, Jones SJM, Birol I. ABySS: a parallel assembler for short read sequence data. Genome Res. 2009 Jun;19(6):1117–23.

44. Grabherr MG, Haas BJ, Yassour M, Levin JZ, Thompson DA, Amit I, et al. Full-length transcriptome assembly from RNA-Seq data without a reference genome. Nat Biotechnol. 2011 Jul;29(7):644–52.

45. Huang X, Madan A. CAP3: A DNA sequence assembly program. Genome Res. 1999 Sep;9(9):868–77.

46. Karim S, Singh P, Ribeiro JMC. A Deep Insight into the Sialotranscriptome of the Gulf Coast Tick, Amblyomma maculatum. PLoS One. 2011 Dec 21;6(12).

47. Nielsen H, Brunak S, Von Heijne G. Machine learning approaches for the prediction of signal peptides and other protein sorting signals. Protein Eng Des Sel. 1999 Jan 1;12(1):3– 9.

48. Kahsay RY, Gao G, Liao L. An improved hidden Markov model for transmembrane protein detection and topology prediction and its applications to complete genomes. Bioinformatics. 2005 May 1;21(9):1853–8.

49. Hansen JE, Lund O, Tolstrup N, Gooley AA, Williams KL, Brunak S. NetOglyc: Prediction of mucin type O-glycosylation sites based on sequence context and surface accessibility. Glycoconjugate J 1998 152. 1998;15(2):115–30.

50. Robinson MD, McCarthy DJ, Smyth GK. edgeR: a Bioconductor package for differential expression analysis of digital gene expression data. Bioinformatics. 2010 Nov 11;26(1):139–40.

51. Ge SX, Son EW, Yao R. iDEP: an integrated web application for differential expression and pathway analysis of RNA-Seq data. BMC Bioinformatics. 2018 Dec 19;19(1).

52. Anacker RL, Mann RE, Gonzales C. Reactivity of monoclonal antibodies to Rickettsia rickettsii with spotted fever and typhus group rickettsiae. J Clin Microbiol. 1987;25(1):167–71.

53. Policastro PF, Munderloh UG, Fischer ER, Hackstadt T. Rickettsia rickettsii growth and temperature-inducible protein expression in embryonic tick cell lines. J Med Microbiol. 1997;46(10):839–45.

54. van Rooijen N, Hendrikx E. Liposomes for specific depletion of macrophages from organs and tissues. Methods Mol Biol. 2010;605:189–203.

55. Lehenkari PP, Kellinsalmi M, Näpänkangas JP, Ylitalo K V., Mönkkönen J, Rogers MJ, et al. Further insight into mechanism of action of clodronate: Inhibition of mitochondrial ADP/ATP translocase by a nonhydrolyzable, adenine-containing metabolite. Mol Pharmacol. 2002 May 1;61(5):1255–62.

56. Kwon H, Mohammed M, Franzén O, Ankarklev J, Smith RC. Single-cell analysis of mosquito hemocytes identifies signatures of immune cell subtypes and cell differentiation. Elife. 2021 Jul 1;10.

57. Aung KM, Boldbaatar D, Umemiya-Shirafuji R, Liao M, Xuenan X, Suzuki H, et al. Scavenger Receptor Mediates Systemic RNA Interference in Ticks. PLoS One. 2011 Dec 1;6(12):e28407.

58. Oliver JD, Dusty Loy J, Parikh G, Bartholomay L. Comparative analysis of hemocyte phagocytosis between six species of arthropods as measured by flow cytometry. J Invertebr Pathol. 2011 Oct 1;108(2):126–30.

59. Kuhn KH. Mitotic activity of the hemocytes in the tick Ixodes ricinus (Acari; Ixodidae). Parasitol Res. 1996;82(6):511–7.

60. Kadota K, Walter S, Claveria FG, Igarashi I, Taylor D, Fujisaki K. Morphological and Populational Characteristics of Hemocytes ofOrnithodoros moubata Nymphs During the Ecdysial Phase. J Med Entomol. 2003 Nov 1;40(6):770–6.

61. Leite THJF, Ferreira AGA, Imler JL, Marques JT. Distinct Roles of Hemocytes at Different Stages of Infection by Dengue and Zika Viruses in Aedes aegypti Mosquitoes. Front Immunol. 2021 May 13;12.

62. Kotsyfakis M, Kopáček P, Franta Z, Pedra JHF, Ribeiro JMC. Deep Sequencing Analysis of the Ixodes ricinus Haemocytome. PLoS Negl Trop Dis. 2015 May 13;9(5).

63. Waltzer L, Gobert V, Osman D, Haenlin M. Transcription factor interplay during Drosophila haematopoiesis. Int J Dev Biol. 2010;54(6–7):1107–15.

64. Liu M, Liu S, Liu H. Recent insights into hematopoiesis in crustaceans. Fish Shellfish Immunol Reports. 2021 Dec 1;2:100040.

65. Lin X, Söderhäll I. Crustacean hematopoiesis and the astakine cytokines. Blood. 2011 Jun 16;117(24):6417–24.

66. Saelee N, Noonin C, Nupan B, Junkunlo K, Phongdara A, Lin X, et al. β-Thymosins and Hemocyte Homeostasis in a Crustacean. PLoS One. 2013 Apr 2;8(4):e60974.

67. Charoensapsri W, Sangsuriya P, Lertwimol T, Gangnonngiw W, Phiwsaiya K, Senapin S. Laminin receptor protein is implicated in hemocyte homeostasis for the whiteleg shrimp Penaeus (Litopenaeus) vannamei. Dev Comp Immunol. 2015 Jul 1;51(1):39–47.

68. Kanost MR, Jiang H. Clip-domain serine proteases as immune factors in insect hemolymph. Curr Opin insect Sci. 2015 Oct 1;11:47–55.

69. Matsushita M, Fujita T. The role of ficolins in innate immunity. Immunobiology. 2002;205(4–5):490–7.

70. Gokudan S, Muta T, Tsuda R, Koori K, Kawahara T, Seki N, et al. Horseshoe crab acetyl group-recognizing lectins involved in innate immunity are structurally related to fibrinogen. Proc Natl Acad Sci U S A. 1999 Aug 8;96(18):10086.

71. Adema CM, Hertel LA, Miller RD, Loker ES. A family of fibrinogen-related proteins that precipitates parasite-derived molecules is produced by an invertebrate after infection. Proc Natl Acad Sci U S A. 1997 Aug 5;94(16):8691–6.

72. Dimopoulos G, Casavant TL, Chang S, Scheetz T, Roberts C, Donohue M, et al. Anopheles gambiae pilot gene discovery project: identification of mosquito innate immunity genes from expressed sequence tags generated from immune-competent cell lines. Proc Natl Acad Sci U S A. 2000 Jun 6;97(12):6619–24.

73. Dimopoulos G, Christophides GK, Meister S, Schultz J, White KP, Barillas-Mury C, et al. Genome expression analysis of Anopheles gambiae: Responses to injury, bacterial challenge, and malaria infection. Proc Natl Acad Sci U S A. 2002 Jun 25;99(13):8814–9.

74. Rego ROM, Hajdušek O, Kovář V, Kopáček P, Grubhoffer L, Hypša V. Molecular cloning and comparative analysis of fibrinogen-related proteins from the soft tick Ornithodoros moubata and the hard tick Ixodes ricinus. Insect Biochem Mol Biol. 2005 Sep 1;35(9):991–1004.

75. Michel T, Relchhart JM, Hoffmann JA, Royet J. Drosophila Toll is activated by Gram-positive bacteria through a circulating peptidoglycan recognition protein. Nature. 2001 Dec 13;414(6865):756–9.

76. Kanagawa M, Satoh T, Ikeda A, Adachi Y, Ohno N, Yamaguchi Y. Structural insights into recognition of triple-helical beta-glucans by an insect fungal receptor. J Biol Chem. 2011 Aug 19;286(33):29158–65.

77. Valanne S, Wang J-H, Rämet M. The Drosophila Toll Signaling Pathway. J Immunol. 2011 Jan 15;186(2):649–56.

78. Wu LP, Anderson K V. Regulated nuclear import of Rel proteins in the Drosophila immune response. Nature. 1998 Mar 5;392(6671):93–7.

79. Lemaitre B, Kromer-Metzger E, Michaut L, Nicolas E, Meister M, Georgel P, et al. A recessive mutation, immune deficiency (imd), defines two distinct control pathways in the Drosophila host defense. Proc Natl Acad Sci U S A. 1995 Oct 10;92(21):9465–9.

80. Lu S, Wang J, Chitsaz F, Derbyshire MK, Geer RC, Gonzales NR, et al. CDD/SPARCLE: the conserved domain database in 2020. Nucleic Acids Res. 2020 Jan 1;48(D1):D265–8.

81. Melcarne C, Ramond E, Dudzic J, Bretscher AJ, Kurucz É, Andó I, et al. Two Nimrod receptors, NimC1 and Eater, synergistically contribute to bacterial phagocytosis in Drosophila melanogaster. FEBS J. 2019 Jul 1;286(14):2670–91.

82. Kurucz É, Márkus R, Zsámboki J, Folkl-Medzihradszky K, Darula Z, Vilmos P, et al. Nimrod, a Putative Phagocytosis Receptor with EGF Repeats in Drosophila Plasmatocytes. Curr Biol. 2007 Apr 3;17(7):649–54.

83. Estévez-Lao TY, Hillyer JF. Involvement of the Anopheles gambiae Nimrod gene family in mosquito immune responses. Insect Biochem Mol Biol. 2014 Jan;44(1):12–22.

84. Bretscher AJ, Honti V, Binggeli O, Burri O, Poidevin M, Kurucz É, et al. The Nimrod transmembrane receptor Eater is required for hemocyte attachment to the sessile compartment in Drosophila melanogaster. Biol Open. 2015 Mar 15;4(3):355–63.

85. Rodrigues J, Brayner FA, Alves LC, Dixit R, Barillas-Mury C. Hemocyte differentiation mediates innate immune memory in Anopheles gambiae mosquitoes. Science (80-). 2010 Sep 10;329(5997):1353–5.

86. Bryant WB, Michel K. Anopheles gambiae hemocytes exhibit transient states of activation. Dev Comp Immunol. 2016 Feb 1;55:119–29.

87. Smith RC, King JG, Tao D, Zeleznik OA, Brando C, Thallinger GG, et al. Molecular profiling of phagocytic immune cells in Anopheles gambiae reveals integral roles for hemocytes in mosquito innate immunity. Mol Cell Proteomics. 2016;15(11):3373–87.

88. Shaw DK, Wang X, Brown LJ, Chávez ASO, Reif KE, Smith AA, et al. Infection-derived lipids elicit an immune deficiency circuit in arthropods. Nat Commun 2017 81. 2017 Feb 14;8(1):1–13.

89. Jordan MB, Van Rooijen N, Izui S, Kappler J, Marrack P. Liposomal clodronate as a novel agent for treating autoimmune hemolytic anemia in a mouse model. Blood. 2003 Jan 15;101(2):594–601.

90. Lemaitre B, Hoffmann J. The Host Defense of Drosophila melanogaster. http://dx.doi.org/101146/annurev.immunol25022106141615. 2007 Mar 21;25:697–743.

91. Blow F, Douglas AE. The hemolymph microbiome of insects. J Insect Physiol. 2019 May 1;115:33–9.

92. Sf H, Burgdorfer W. Interactions between rickettsial endosytobionts and their tick hosts. In: Selwemmler W, Gassner G, editors Insect Endocytobiosis: Morphology, Physiology, Genetics, Evolution. Boca Raton, Florida: CRC Press; 1989. p. 235–51.

93. Baldridge GD, Kurtti TJ, Burkhardt N, Baldridge AS, Nelson CM, Oliva AS, et al. Infection of Ixodes scapularis ticks with Rickettsia monacensis expressing green fluorescent protein: A model system. J Invertebr Pathol. 2007 Mar;94(3):163.

94. Harris EK, Jirakanwisal K, Verhoeve VI, Fongsaran C, Suwanbongkot C, Welch MD, et al. Role of Sca2 and RickA in the Dissemination of Rickettsia parkeri in Amblyomma maculatum. Infect Immun. 2018 Jun 1;86(6).

95. Garcia Guizzo M, Budachetri K, Adegoke A, Ribeiro JM, Karim S. Rickettsia parkeri infection modulates the sialome and ovariome of the Gulf Coast tick, Amblyomma maculatum. Front Microbiol. 1AD;0:4233.

96. Harris EK, Verhoeve VI, Banajee KH, Macaluso JA, Azad AF, Macaluso KR. Comparative vertical transmission of Rickettsia by Dermacentor variabilis and Amblyomma maculatum. Ticks Tick Borne Dis. 2017 Jun 1;8(4):598.

97. Wright CL, Gaff HD, Sonenshine DE, Hynes WL. Experimental vertical transmission of Rickettsia parkeri in the Gulf Coast tick, Amblyomma maculatum. Ticks Tick Borne Dis. 2015 Jul 1;6(5):568–73.

98. Lee JK, Moraru GM, Stokes J V., Benton AN, Wills RW, Nabors HP, et al. Amblyomma maculatum-associated rickettsiae in vector tissues and vertebrate hosts during tick feeding. Exp Appl Acarol. 2019 Feb 1;77(2):187.

99. Saito TB, Bechelli J, Smalley C, Karim S, Walker DH. Vector Tick Transmission Model of Spotted Fever Rickettsiosis. Am J Pathol. 2019;189(1):115–23.

100. King JG, Hillyer JF. Spatial and temporal in vivo analysis of circulating and sessile immune cells in mosquitoes: Hemocyte mitosis following infection. BMC Biol. 2013;11.

101. Leitão AB, Sucena É. Drosophila sessile hemocyte clusters are true hematopoietic tissues that regulate larval blood cell differentiation. Elife. 2015 Apr 2;2015(4):1–38.

102. Speck NA, Gilliland DG. Core-binding factors in haematopoiesis and leukaemia. Nat Rev Cancer 2002 27. 2002;2(7):502–13.

103. Fossett N, Hyman K, Gajewski K, Orkin SH, Schulz RA. Combinatorial interactions of Serpent, Lozenge, and U-shaped regulate crystal cell lineage commitment during Drosophila hematopoiesis. Proc Natl Acad Sci U S A. 2003 Sep 9;100(20):11451.

104. Waltzer L, Ferjoux G, Bataillé L, Haenlin M. Cooperation between the GATA and RUNX factors Serpent and Lozenge during Drosophila hematopoiesis. EMBO J. 2003 Dec 12;22(24):6516.

105. Rehorn KP, Thelen H, Michelson AM, Reuter R. A molecular aspect of hematopoiesis and endoderm development common to vertebrates and Drosophila. Development. 1996 Dec 1;122(12):4023–31.

106. Lebestky T, Chang T, Hartenstein V, Banerjee U. Specification of Drosophila hematopoietic lineage by conserved transcription factors. Science (80-). 2000 Apr 7;288(5463):146–9.

107. Umemiya-Shirafuji R, Boldbaatar D, Liao M, Battur B, Rahman MM, Kuboki T, et al. Target of rapamycin (TOR) controls vitellogenesis via activation of the S6 kinase in the fat body of the tick, Haemaphysalis longicornis. Int J Parasitol. 2012 Oct;42(11):991–8.

108. Boldbaatar D, Battur B, Umemiya-Shirafuji R, Liao M, Tanaka T, Fujisaki K. GATA transcription, translation and regulation in Haemaphysalis longicornis tick: analysis of the cDNA and an essential role for vitellogenesis. Insect Biochem Mol Biol. 2010 Jan;40(1):49–57.

109. Kuniyori M, Sato N, Yokoyama N, Kawazu S, Xuan X, Suzuki H, et al. Vitellogenin-2 Accumulation in the Fat Body and Hemolymph of Babesia-Infected Haemaphysalis longicornis Ticks. Front Cell Infect Microbiol. 2022 Jun 21;12:1.

110. Boquet I, Boujemaa R, Carlier MF, Préat T. Ciboulot regulates actin assembly during Drosophila brain metamorphosis. Cell. 2000 Sep 15;102(6):797–808.

111. Herrmann D, Hatta M, Hoffmeister-Ullerich SAH. Thypedin, the multi copy precursor for the hydra peptide pedin, is a beta-thymosin repeat-like domain containing protein. Mech Dev. 2005 Nov;122(11):1183–93.

112. Sunyakumthorn P, Petchampai N, Grasperge BJ, Kearney MT, Sonenshine DE, Macaluso KR. Gene expression of tissue-specific molecules in ex vivo Dermacentor variabilis (Acari: Ixodidae) during rickettsial exposure. J Med Entomol. 2013 Sep;50(5):1089–96.

113. Belkin AM. Extracellular TG2: emerging functions and regulation. FEBS J. 2011 Dec 1;278(24):4704–16.

114. Lin X, Söderhäll K, Söderhäll I. Transglutaminase activity in the hematopoietic tissue of a crustacean, Pacifastacus leniusculus, importance in hemocyte homeostasis. BMC Immunol. 2008 Oct 7;9(1):1–11.

115. Junkunlo K, Söderhäll K, Söderhäll I. Transglutaminase inhibition stimulates hematopoiesis and reduces aggressive behavior of crayfish, Pacifastacus leniusculus. J Biol Chem. 2019 Jan 11;294(2):708–15.

116. Junkunlo K, Söderhäll K, Söderhäll I. Clotting protein – An extracellular matrix (ECM) protein involved in crustacean hematopoiesis. Dev Comp Immunol. 2018 Jan 1;78:132– 40.

117. Lin X, Novotny M, Söderhäll K, Söderhäll I. Ancient Cytokines, the Role of Astakines as Hematopoietic Growth Factors. J Biol Chem. 2010 Sep 10;285(37):28577–86.

118. Rosa RD, Capelli-Peixoto J, Mesquita RD, Kalil SP, Pohl PC, Braz GR, et al. Exploring the immune signalling pathway-related genes of the cattle tick Rhipicephalus microplus: From molecular characterization to transcriptional profile upon microbial challenge. Dev Comp Immunol. 2016 Jun 1;59:1–14.

119. Liu XY, Bonnet SI. Hard tick factors implicated in pathogen transmission. PLoS Negl Trop Dis. 2014;8(1):5.

120. McClure Carroll EE, Wang X, Shaw DK, O’Neal AJ, Oliva Chávez AS, Brown LJ, et al. p47 licenses activation of the immune deficiency pathway in the tick Ixodes scapularis. Proc Natl Acad Sci U S A. 2019 Jan 2;116(1):205–10.

121. Jo Caimano M, Anguita J, bioGUNE C, Job Lopez SE, Kleino A, Oliva Chávez AS, et al. Tick Humoral Responses: Marching to the Beat of a Different Drummer. 2017;

122. Smith AA, Pal U. Immunity-related genes in Ixodes scapularis-perspectives from genome information. Front Cell Infect Microbiol. 2014;4(AUG).

123. Palmer WJ, Jiggins FM. Comparative Genomics Reveals the Origins and Diversity of Arthropod Immune Systems. Mol Biol Evol. 2015 Aug 1;32(8):2111.

124. Sigle LT, Hillyer JF. Eater and draper are involved in the periostial haemocyte immune response in the mosquito Anopheles gambiae. Insect Mol Biol. 2018 Aug 1;27(4):429–38.

125. Buresova V, Hajdusek O, Franta Z, Sojka D, Kopacek P. IrAM-An alpha2-macroglobulin from the hard tick Ixodes ricinus: characterization and function in phagocytosis of a potential pathogen Chryseobacterium indologenes. Dev Comp Immunol. 2009 Apr;33(4):489–98.

126. Buresova V, Hajdusek O, Franta Z, Loosova G, Grunclova L, Levashina EA, et al. Functional genomics of tick thioester-containing proteins reveal the ancient origin of the complement system. J Innate Immun. 2011 Oct;3(6):623–30.

